# Characterisation of transgenic lines labelling reticulospinal neurons in larval zebrafish

**DOI:** 10.1101/2024.12.20.629714

**Authors:** Elena M.D. Collins, Pedro T.M. Silva, Aaron D. Ostrovsky, Sabine L. Renninger, Ana R. Tomás, Ruth Diez del Corral, Michael B. Orger

**Affiliations:** Champalimaud Research, Champalimaud Foundation; International Neuroscience Doctoral Program, Champalimaud Foundation

**Keywords:** control of locomotion, reticulospinal circuitry, transgene expression, zebrafish

## Abstract

From lamprey to monkeys, the organization of the descending control of locomotion is conserved across vertebrates. Reticulospinal neurons (**RSNs**) form a bottleneck for descending commands, receiving innervation from diencephalic and mesencephalic locomotor centres and providing locomotor drive to spinal motor circuits. Given their optical accessibility in early development, larval zebrafish offer a unique opportunity to study reticulospinal circuitry. In fish, RSNs are few, highly stereotyped, uniquely identifiable, large neurons spanning from the midbrain to the medulla. Classically labelled by tracer dye injections into the spinal cord, recent advances in genetic tools have facilitated the targeted expression of transgenes in diverse brainstem neurons of larval zebrafish. Here, we provide a comparative characterization of four existing and three newly established transgenic lines in larval zebrafish. We determine which identified neurons are consistently labelled and offer projection-specific genetic access to subpopulations of RSNs. We showcase transgenic lines that label most or all RSNs (*nefma, adcyap1b^ccu96Et^*) or subsets of RSNs, including ipsilateral (*vsx2, calca^ccu75Et^*), contralateral (*pcp4a^ccu97Tg^*) or all (*tiam2a^y264Et^*) components of the Mauthner array, or midbrain-only RSNs (*s1171tEt*). In addition to RSNs, selected transgenic lines *(nefma, s1171tEt, calca^ccu75Et^)* labelled other potential neurons of interest in the brainstem. For those, we performed *in situ* hybridisation to show expression patterns of several excitatory and inhibitory neurotransmitters at larval stages as well as glutamatergic expression patterns in juvenile fish. We provide an overview of transgene expression in the brainstem of larval zebrafish that serves to lay a foundation for future studies in the supraspinal control of locomotion.

**Significance Statement:** Genetic access to subpopulations of brainstem neurons greatly facilitates the dissection of supraspinal circuitry and function. Here, we present several new transgenic lines and rigorously describe existing ones, all in terms of their degree of overlap with the reticulospinal system, variability in transgenic labelling and neurotransmitter identity. Having transgenic access to different subpopulations of reticulospinal neurons enables targeted functional calcium imaging, anatomical tracing and optogenetic manipulations to decipher the role of individual reticulospinal neurons in movement production, ultimately clarifying existing understanding and facilitating future studies in the supraspinal control of locomotion.

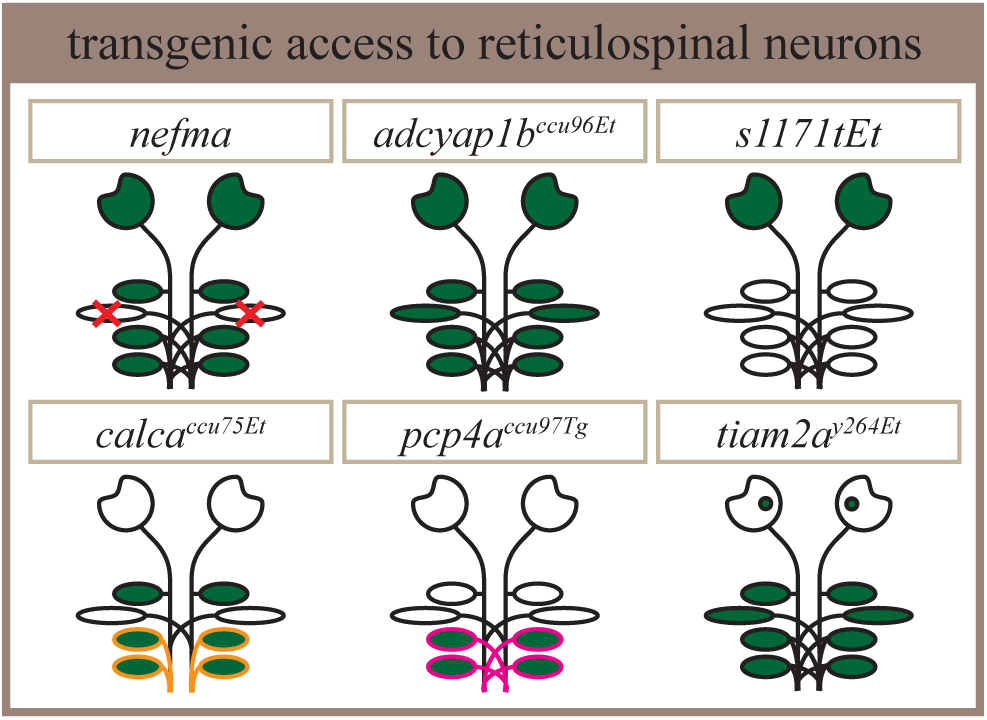

## Introduction

Animals are constantly faced with the need to produce flexible behaviour in response to a rapidly changing environment. From foraging for food, finding a mate, to escaping from predators – movement is central to survival. As such, many regions of the central nervous system are dedicated to its production (Brownstone and Chopek, 2018). A key processing site for descending locomotor commands is the reticular formation (**RF**), situated in the brain-stem and comprised of several nuclei. The RF contains cholinergic (Le Ray et al., 2022; Josset et al., 2018), monoaminergic (McLean and Fetcho, 2004), GABA/glycinergic and glutamatergic neurons (Higashijima et al., 2004), with glutamatergic reticulospinal neurons (**RSNs**) forming the key excitatory descending output (Perreault and Giorgi, 2019). RSNs receive descending input from the mesencephalic and diencephalic locomotor regions (**MLR, DLR**) and the cerebellum, as well as ascending signals from the spinal cord allowing for sensorimotor integration. Their main output is to central pattern generators comprised of spinal inter- and motor neurons, responsible for movement production (for a comprehensive review on the descending control of locomotion see (Grillner and El Manira, 2020)). A central question in neuroscience has long been whether different behaviours are generated by distinct or overlapping populations of RSNs (Siegel, 1979), and if so, whether activity is localised or distributed across the brainstem. The ability to label, activate and manipulate subsets of RSNs is critical in addressing these questions.

Larval zebrafish are ideally suited to studying brainstem circuitry due to their small size, high fecundity, array of available genetic tools, optical accessibility and well-characterised behavioural repertoire (Fetcho and Liu, 1998). For decades, researchers have labelled reticulospinal neurons by spinal injections of tracer compounds, such as horseradish peroxidase or dextran-conjugated dyes (Metcalfe et al., 1986; Kimmel et al., 1982; Severi et al., 2014; Orger et al., 2008; Kimmel et al., 1990). While these have been instrumental in disentangling reticulospinal circuity, labelling can be variable depending on the skill of the experimenter, and it is challenging to consistently label descending neurons of small axon calibre. In addition, spinal injections are necessarily damaging to spinal axon tracts and carry the risk of affecting subsequent behavioural studies. A solution comes from several recently established transgenic lines with expression in the brainstem (Eschstruth et al., 2020; Scott and Baier, 2009; Kimura et al., 2006; Marquart et al., 2015). Given the sometimes stochastic expression patterns in Gal4 lines (Akitake et al., 2011), it would be useful to understand precisely which RSNs are labelled by each transgenic line across a number of animals. Additionally, gaining genetic access to all as well as to selected subpopulations of RSNs, for instance ipsi- vs contralaterally-projecting RSNs, would offer an excellent opportunity when combined with optogenetic tools to decipher cell-specific functions in the control of locomotion.

Here, we characterised seven transgenic lines driving gene expression in the brainstem in larval zebrafish at 6 days-post-fertilisation: four existing transgenic lines *(nefma, vsx2, s1171tEt, tiam2a^y264Et^*) and three newly established transgenic lines *(calca^ccu75Et^, pcp4a^ccu97Tg^, adcyap1b^ccu96Et^)*. For each transgenic line, we performed retrograde labelling of reticulospinal neurons followed by immunohistochemistry and quantified overlap across fish at the single-cell level. We showcase transgenic lines that label most or all RSNs (*nefma, adcyap1b^ccu96Et^*), midbrain-only RSNs (*s1171tEt*), or subsets of RSNs, including ipsilateral (*vsx2, calca^ccu75Et^*), contralateral (*pcp4a^ccu97Tg^*) or all (*tiam2a^y264Et^*) components of the Mauthner array.

For select transgenic lines *(nefma, s1171tEt, calca^ccu75Et^)* we performed *in situ* hybridisation to show expression patterns of genes associated with several neurotransmitter phenotypes *(vglut1, vglut2a, vglut2b, chata, gad1b, gad2, glyt1, glyt2)* at larval stages as well as glutamatergic expression in juvenile fish demonstrating consistency across development. By providing a comprehensive overview of transgene expression in the brainstem of larval zebrafish we set a foundation for future studies in the supraspinal control of locomotion.

## Materials and Methods

### Fish husbandry

Adult fish were raised and bred at 28*^◦^*C on a 14h light / 10h dark cycle following standard husbandry methods as detailed in (Martins et al., 2016). All fish colonies were maintained under meticulous plans involving importation of wild types every 1-3 years and line-specific breeding schemes designed to reduce inbreeding depression (Martins et al., 2016). Embryos were collected and larvae were raised at 28*^◦^*C in E3 embryo medium (5 mM NaCl, 0.17 mM KCl, 0.33 mM CaCl2 and 0.33 mM MgSO4, changed daily) at a density of 100 larvae per 200mL until euthanasia at 6 days-post-fertilisation (**dpf**). A subset of fish from the *calca^ccu75Et^* line used for Fig. 1-1 were raised in the presence of the pigmentation inhibitor PTU (1-Phenyl-2-thiourea (PTU) (Sigma-Aldrich), P7629) at 0.2 mM from 1 dpf onwards. From 5 dpf onwards, approximately 10mL of a live L-type rotifer poly-culture (containing 1000-2000 rotifers per mL) were added to each dish twice a day and larvae were allowed to feed freely. Zebrafish do not sexually differentiate until approximately 3 months of age, therefore the sex of the animals cannot be reported. All experimental procedures were approved by the Champalimaud Foundation Ethics Committee and the Portuguese Direcção Geral Veterinária, and were performed according to the European Directive 2010/63/EU.

### Transgenic lines

The following transgenic lines were used in a nacre (mitfa -/-) background (see Table 1). With the exception of *vsx2*, which we received as an mRFP line, and *s1171tEt*, which was crossed with Tg[10xUAS:GCaMP6f^ccu1Tg^] (Van Opbergen et al., 2018), the lines used in the experiments were created by crossing with Tg[10xUAS:GCaMP6fEF05^ccu2Tg^] (Félix et al., 2024).

**Table 1:** Transgenic fish lines used in this study.

| Short name | Transgenic line | Source |
| --- | --- | --- |
| *nefma* | Tg[nefma:KalTa4] | (Eschstruth et al., 2020) |
| *tiam2a*<sup>y264Et</sup> | Tg[tiam2a<sup>y264Et(B)</sup>] | (Marquart et al., 2015) |
| *s1171tEt* | Tg[-0.6hsp70l:Gal4-VP16<sup>s1171tEt+</sup>] | (Thiele et al., 2014; Scott and Baier, 2009) |
| *vsx2* | TgBAC[vsx2:Gal4FF<sup>nns18Tg</sup>] | (Kimura et al., 2013) |
| *calca*<sup>ccu75Et</sup> | Tg[-5.0calca:Gal4FF<sup>ccu75Et</sup>] | this paper |
| *adcyap1b*<sup>ccu96Et</sup> | Tg[-1.7adcyap1b:Gal4FF<sup>ccu96Et</sup>] | this paper |
| *pcp4a*<sup>ccu97Tg</sup> | TgBAC[pcp4a:Gal4FF<sup>ccu97Tg</sup>] | this paper |
|  | Tg[10xUAS:GCaMP6f<sup>ccu1Tg</sup>] | (Van Opbergen et al., 2018) |
|  | Tg[10xUAS:GCaMP6fEF05<sup>ccu2Tg</sup>] | (Félix et al., 2024) |
|  | Tg[UAS:mRFP] | (Asakawa et al., 2008) |

### Cloning

#### adcyap1b:Gal4FF

A 1679bp promoter region upstream of adcyap1b start codon was cloned into pCR™8/GW/ TOPO™ (Invitrogen™) using the following primers:

- 5’-GAACTGGAACACTTGGTGGCAGTATTG-3’
- 5’-GATCTGGCCAGGCTGTAAAGATACAAGAAAG-3’.

The adcyap1b promoter (Entry Clone) was then recombined into an Gal4FF destination vector (Gateway™ LR recombination, Invitrogen™), derived from the Tol2Kit (Kwan et al., 2007), so that the construct was bracketed by two Tol2 (Kawakami et al., 2000) inverted terminal repeats.

#### calca:Gal4FF

A 5023bp promoter region upstream of calca start codon was cloned into pCR™8/GW/TOPO™ (Invitrogen™) using the following primers:

- 5’-GTGCCTGCTGAGGAGCATAAC-3’
- 5’-GGTCCCCTGTAGTAAAACATC-3’

The calca promoter (Entry Clone) was then recombined into the Tol2 Gal4FF destination vector (Gateway™ LR recombination, Invitrogen™).

#### pcp4a:Gal4FF

The Tg[pcp4a:Gal4FF^ccu97Tg^ line was generated using bacterial artificial chromosome (BAC) recombineering following (Suster et al., 2011). In short, the iTol2-amp cassette was introduced into BAC CH211-231M12. Positive clones were selected to further introduce the Gal4FF-pA-FRT-kan-FRT at the ATG site of *pcp4a*. To do so, homology arms, short DNA sequences of about 300bp that flanked the ATG site and with sequence overlap to the Gal4FF-pA-FRT-kan-FRT, were amplified from the BAC:

- *pcp4a-HI-for* CACACACCAATGCATACATCAAAGCG
- *pcp4a-HI-rev* ACAGTAGCTTCATGGTGGCGCTGGATGAAGAGTATGAAGATGAAGGAA GAAG
- *pcp4a-HII-for* CCAGCCTACACGCGGGTGAGCTTTCCTCCATACACATTGCACA
- *pcp4a-HII-rev* GCGCACATACAATATCCTCCATCCCT

Similarly, the Gal4FF-pA-FRT-kan-FRT cassette was amplified using primers overlapping with the homology arms:

- *pcp4a-GFF-for* CATCTTCATACTCTTCATCCAGCGCCACCATGAAGCTACTGTCTTCTATC-GAAC
- *pcp4a-GFF-rev* TGTGTATGGAGGAAAGCTCACCCGCGTGTAGGCTGGAGCTGCTTC

The 3 DNA fragments were fused using the Gibson Assembly Cloning kit (New England Bio-labs) and subcloned into the pCR2.1-TOPO vector (Invitrogen). The pcp4a-HI-for and pcp4a-HII-rev primer were used for further amplification of the cassette. 500ng of it was used for recombination with the CH211-231M12/iTol2-amp BAC.

#### Line establishment

DNA constructs were injected together with Tol2 transposase mRNA and non-integrating UAS:GFP plasmid into 1-2 cell stage mitfa-/- eggs. Embryos with GFP expression were raised and screened as adults for germ line transmission. Progeny of positive animals with stable expression pattern were selected as founders for the respective Gal4FF driver line.

### Screening

Larvae were pre-screened at 3-4 dpf to select fish with positive expression of GCaMP or RFP. While this is standard practice, we note that some lines appear to have less than Mendelian numbers of offspring with expression presumably due to silencing (Akitake et al., 2011). For instance, the *pcp4a^ccu97Tg^* line seems to be particularly prone to silencing, which can be remedied by setting multiple crosses and rigorous pre-screening.

### Immunohistochemistry

The staining was performed in all seven transgenic lines (Table 1) according to a modified protocol (Randlett et al., 2015).

#### Dye injections

Larvae were raised in standard conditions (see above) until 5 dpf, fed with rotifers in the morning and injected in the afternoon. Larvae were injected while mounted side-ways on a small petri dish containing a layer of agarose gel (SeaKem LE Agarose, #50004, Lonza), following previously described methods (O’Malley et al., 1996). The dye (10,000MW at 50mg/mL of Dextran, Alexa Fluor 647, Invitrogen by Life Technologies) was pressure-injected into the spinal cord near myomere 8 using a pulled capillary glass needle (GC100F-10, Harvard Apparatus) positioned with a micro-manipulator (MN-153, Narishige) and a stereo-microscope (Stereo Discovery V8, Zeiss). Pressure was applied using a pneumatic picopump (WPI, PV820). Following injection, larvae were allowed to recover overnight and checked for normal swimming behaviour before euthanasia.

#### Immunohistochemistry

At 6 dpf, larvae were anaesthetized in 15mM tricaine (E10521, Sigma-Aldrich) for 10 min and fixed with 4% paraformaldehye (**PFA**) for 2 hours at room temperature while covered and with agitation. From this step onwards, larvae were kept in darkness. To stop fixation, larvae were rinsed and washed 2 x 5 min in phosphate-buffered saline with 0.25% Triton (**PBT**) with agitation. For epitope retrieval, larvae were rinsed twice, washed 1 x 5 min, and finally incubated in 150mM Tris-HCl with pH 9.0 at 70*^◦^*C in a water bath. Following incubation, samples were cooled on ice and washed 3 x 5 min in PBT. To permeabilise, larvae were incubated in 0.05% Trypsin-EDTA in phosphate-buffered saline (**PBS**) for 5 min on ice, followed by a rinse and 2 x 5 min washes in PBT. Next, larvae were incubated in blocking solution (PBS, bovine serum albumin, normal goat serum, dimethyl sulfoxide, Triton, azide, sterilised *H~*_2_O) at 4*^◦^*C overnight. A primary antibody solution was prepared by diluting primary antibodies for anti-tERK and anti-GFP or anti-mCherry (Table 2) in blocking solution at 1:500. Blocking solution was replaced by the primary antibody solution and larvae were allowed to incubate for at least three over-nights at 4*^◦^*C under cover with agitation. To remove primary antibodies, larvae were rinsed three times and washed 3 x 30 min in PBT. A secondary antibody solution was prepared by diluting secondary antibodies in blocking solution at 1:500. Larvae were incubated in secondary antibody solution for at least three over-nights at 4*^◦^*C with agitation. To remove secondary antibodies (Table 2), larvae were rinsed three times and washed 3 x 30 min in PBT. Samples were stored in PBT at 4*^◦^*C in the dark until imaging.

**Table 2:** Reagents used for immunohistochemistry experiments in this study.

| Reagents | Source | Identifier |
| --- | --- | --- |
| Dye for Backfills |  |  |
| Dextran, Alexa Fluor™ 647 | Life Technologies | D22914 |
| Primary Antibodies |  |  |
| Chicken anti-GFP | Abcam Limited | ABCAM-13970 |
| Mouse anti-tERK | Cell Signaling Technology | #4696 |
| Rabbit anti-mCherry | Abcam Limited | ABCAM-167453 |
| Secondary Antibodies |  |  |
| Goat anti-Chicken IgY Alexa Fluor™ 488 | Life Technologies | A-11039 |
| Goat anti-Mouse IgG Alexa Fluor™ 568 | Life Technologies | A-11004 |
| Goat anti-Mouse IgG Alexa Fluor™ 488 | Life Technologies | A-11001 |
| Goat anti-Rabbit IgG Alexa Fluor™ 568 | Life Technologies | A-11011 |

### *In situ* hybridisation chain reaction in larval zebrafish

*In situ* hybridisation chain reaction (**isHCR**) reagents, including probes, hairpins, and buffers, were purchased from Molecular Instruments (Los Angeles, CA, USA) for detection of mRNAs. The staining was performed according to the “HCR v3.0 protocol for whole-mount zebrafish embryos and larvae” protocol provided by Molecular Instruments (Choi et al., 2018). The following transgenic lines were used in a nacre (mitfa -/-) background: Tg[nefma:KalTa4, 10xUAS:GCaMP6fEF05], Tg[-5.0calca:Gal4FF^ccu75Et^, 10xUAS:GCaMP6fEF05] and Tg[-0.6hsp70l:Gal4-VP16^s1171tEt+^, 10xUAS:GCaMP6f].

#### Tissue fixation

At 6 dpf, larvae were anaesthetised in 15mM tricaine (E10521, Sigma-Aldrich) and fixed with 4% PFA overnight at 4*^◦^*C while covered and gently agitated. From this step onwards, larvae were continuously kept in darkness.

#### Tissue preparation

On the following day, larvae were washed 3 x 5 min in PBS to stop fixation, dehydrated and permeabilised with a series of 100% methanol (**MeOH**) washes and stored at -20*^◦^*C for up to 6 months before use. MeOH-fixed larvae were re-hydrated progressively with a series of graded MeOH in PBS with Tween 0.1% (**PBST**) washes at room temperature with agitation for 5 min each. Next, larvae were treated with proteinase K (30*µ*g/mL) for 45 minutes at room temperature, rinsed twice with PBST, post-fixed with 4% PFA for 20 minutes at room temperature and finally washed thoroughly 5 x 5 min with PBST with agitation.

#### Detection

Larvae were pre-hybridised with 500*µ*L of probe hybridisation buffer for 30 minutes at 37*^◦^*C. Probe solutions were prepared by adding 2pmols of each probe set (Table 3) to 500*µ*L of probe hybridisation buffer at 37*^◦^*C. The pre-hybridisation solution was replaced with the probe solution mix and larvae were incubated overnight at 37*^◦^*C. The following day, excess probes were removed by washing larvae 4 x 15 min with 500*µ*L of probe wash buffer at 37*^◦^*C, followed by 2 x 5 min washes with 5x sodium chloride sodium citrate buffered with 0.1% Tween (**5xSSCT**) at room temperate with agitation. Note that in the isHCR 3.0 method, detection of an mRNA requires a mixture of primers, each of them including a part that is complementary to the target RNA and another part that is used for amplification of the signal (named B1 to B5). Table 3 summarises the probes indicating their target and the amplification reagent used.

**Table 3:** Reagents used for *in situ* hybridisation experiments in this study.

| Reagents | Source | Identifier |
| --- | --- | --- |
| HCR probes |  |  |
| gad1b-B1: PRA299 | Molecular Instruments | NM194419.1 |
| gad2-B1: PRH743 | Molecular Instruments | NM001017708.2 |
| vglut2.1-B2: PRB128 | Molecular Instruments | NM001128821.1 |
| vglut2.2-B2: PRG620 | Molecular Instruments | NM001009982.1 |
| chATa-B3: RTA359 | Molecular Instruments | NM001130719 |
| glyt1-B4 (slc6a9): RTM844 | Molecular Instruments | NM001030073 |
| glyt2-B4 (slc6a5): B4RTO773 | Molecular Instruments | NM001009557.1 |
| vglut1-B5 (slc17a7a): RTM843 | Molecular Instruments | NM001098755 |

#### Amplification

Pre-amplification was performed by incubating with 500*µ*L of amplification buffer for 30 min at room temperature. Next, 30pmol of hairpin h1 and 30pmol of hairpin h2 were prepared by snap-cooling 10*µ*L of 3*µ*M stock: hairpins were heated separately at 95*^◦^*C for 90 seconds and allowed to cool down to room temperature in a dark drawer for 30 min. After cooling down, the hairpin solution was prepared by adding snap-cooled h1 and h2 hairpins to 500*µ*L of amplification buffer at room temperature. The pre-amplification solution was replaced with the hairpin solution and larvae were incubated overnight in the dark at room temperature. On the next day, excess hairpins were removed by washing the samples with 500*µ*L of 5xSSCT at room temperature with agitation for 2 x 5min, 2 x 30min and 1 x 5min. Samples were stored at 4*^◦^*C in 5xSSCT in the dark until imaging.

### *In situ* hybridisation chain reaction in juvenile zebrafish

To perform *in situ* hybridisation chain reaction in juvenile fish, the aforementioned *isHCR* 3.0 protocol for larval zebrafish from Molecular Instruments (Los Angeles, CA) was adapted using methods from (Kenney et al., 2021) as well as considering advice from Molecular Instruments on adapting the protocol for adult zebrafish (Dr. Chanpreet Singh, personal communication).

#### Tissue fixation

Juvenile zebrafish Tg[-0.6hsp70l:Gal4-VP16^s1171tEt+^] at 4 weeks post-fertilisation were anaesthetised in 15mM tricaine tricaine (E10521, Sigma-Aldrich) for 5 minutes and immersed in ice-cold fish facility water for 20 minutes. Absence of reflexes were assessed before fish were dissected caudal to cloaca using a razor blade and heads were placed in ice-cold PBS for 10 minutes to let blood drain. Samples were fixed in 4% PFA overnight at 4*^◦^*C with gentle agitation.

#### Tissue preparation

The next day, samples were washed 2x for 5min in cold PBS, followed by careful dissection of brains into cold, sterile PBS and stored at 4*^◦^*C until further processing. Following dissection, samples were split into groups of 4 and washed 3x for 5min in PBS at room temperature with gentle agitation. Then, samples were dehydrated using a series of MeOH/PBS mixtures (1h each in 20%, 40%, 60%, 80% and 2x 100% MeOH). Samples were washed 2x for 5min in 100% MeOH and incubated in 5% hydrogen peroxide in MeOH overnight at 4*^◦^*C. The next day, samples were rehydrated using a series of MeOH/PBS mixtures (1h each in 80%, 60%, 40% and 20% MeOH). Samples were washed 2x for 5min in PBS and then 2x for 1h in PBS with 0.2% TritonX-100 (**PTx.2**). Samples were then washed overnight at 37*^◦^*C in permeabilisation solution (PTx.2, 0.3M glycine, 20% DMSO).

#### Detection and amplification

The next day, samples were washed 2x for 5min in PTx.2. From here on, RNA detection was performed following the Detection and Amplification sections described above, with the following modifications: In the detection stage, probe solutions were prepared by adding 10pmols of each probe set to 500*µ*L of probe hybridisation buffer at 37*^◦^*C. In the Amplification stage, 45pmol of hairpin h1 and 30pmol of hairpin h2 were used by snap-cooling 15*µ*L of 3*µ*M stock.

#### Clearing

Samples were washed 2x for 5min in 50%SSC/50%PBS, dehydrated using a series of MeOH/PBS mixtures (1h each in 20%, 40%, 60%, 80% and 2x 100% MeOH) and incubated in 100% MeOH overnight at 4*^◦^*C. The following day, samples were incubated in 66% dichloromethane in MeOH for 3h at room temperature. Samples were washed 2x for 15min dichloromethane and then stored in dibenzyl ether until imaging.

### Imaging and image processing

#### Imaging conditions for larval zebrafish

Samples were cut to remove most of the spinal cord for easier handling and mounted dorsal-side up in 1% low-melting point agarose (Ultra-Pure LMP Agarose, Cat#16520100, Invitrogen by Life Technologies) prepared in PBS or SSC. Imaging of samples was performed on an upright confocal laser-scanning microscope (Zeiss, LSM 980) with a 25x multi-immersion objective (Zeiss, NA 0.8, Plan-Apochromat), using laser wavelengths 488, 594 and 650 nm. Images were acquired with a pixel size of 0.79*µ*m in x and y and sampled with a 1*µ*m interval in the z axis. Pinhole size was 35*µ*m, corresponding to 1.9*µ*m confocal section.

#### Imaging conditions for juvenile zebrafish

Samples were mounted dorsal-side up by attaching to a small petri dish lid using UV-cured glue (Bondic, UV Liquid Plastic Welder Starter Kit) and filling the lid with immersion oil (Gelest, PHENYLMETHYLSILOXANE OLIGOMER #PDM-7040) matching the refractive index of dibenzyl ether. Imaging of samples was performed on an upright confocal laser-scanning microscope (Zeiss, LSM 980) with a 25x multi-immersion objective (Zeiss, NA 0.8, Plan-Apochromat), using laser wavelengths 594 and 650 nm. Images were acquired with a pixel size of 0.79*µ*m in x and y and sampled with a 2*µ*m interval in the z axis. Pinhole size was 35*µ*m, corresponding to 1.9*µ*m confocal section.

#### Image processing

Image analysis was performed using custom-written scripts in ImageJ processing package (Fiji). Briefly, where applicable, tiles were stitched and converted to a resolution that matches an in-house reference image. Using ANTs (Avants et al., 2009), each fish was registered to an in-house reference image of total extracellular signal-related kinase (**tERK**), which labels all neurons, or a transgenic line-specific reference image. All image processing was performed on a computer with 64.0GB RAM and a Intel(R) Core(TM) i7-5820K CPU @ 3.30GHz 3.30 GHz processor.

### Data Availability

Stacks of example fish for each transgenic line were uploaded in a data sharing repository (www.zenodo.org, doi: 10.5281/zenodo.1510272). First channel (green) is the transgenic line, second channel (grey) is the tERK counter-stain, third channel (magenta) is the reticulospinal backfill.

## Results

In this study we characterised four existing (*nefma*, Tg[nefma:KalTa4]; *tiam2a^y264Et^*, Tg[tiam2a^y264Et(B)^]; *s1171tEt*, Tg[-0.6hsp70l:Gal4-VP16^s1171tEt+^]; *vsx2*, TgBAC[vsx2: Gal4FF^nns18Tg^]) and three new (*calca^ccu75Et^*, Tg[-5.0calca: Gal4FF^ccu75Et^]; *pcp4a^ccu97Tg^*, TgBAC[pcp4a:Gal4FF^ccu97Tg^]; *adcyap1b^ccu96Et^*, Tg[-1.7adcyap1b:Gal4FF^ccu96Et^]) transgenic lines. Each line was crossed with Tg[10xUAS:GCaMP6fEF05] unless reported otherwise. We note that transgenic lines based on small promoter regions will have expression patterns that may reflect the locus of insertion in the genome. We have therefore denoted these lines as enhancer traps, indicating that they are not expected to be faithful reporters of the natural expression pattern of the genes from which the construct was derived. Here we aim to characterise these transgenic lines as tools in themselves, to lay a basis for dissecting the role of premotor brainstem neurons in movement production. We chose these transgenic lines as they label reticulospinal (**RSNs**) and other potential neurons of interest in the brainstem of larval zebrafish at 6 days-post-fertilisation (**dpf**) (Fig. 1A). Briefly, *nefma*, *adcyap1b^ccu96Et^* and *tiam2a^y264Et^* label cells across the brainstem, *s1171tEt* in the tegmentum, and *pcp4a^ccu97Tg^*, *calca^ccu75Et^* and *vsx2* in the hindbrain. Larval zebrafish of each transgenic line were injected with dextran-conjugated tracer dye to retrogradely label RSNs and compared to their transgene expression profiles following immunohistochemistry and confocal imaging.

**Figure 1:**
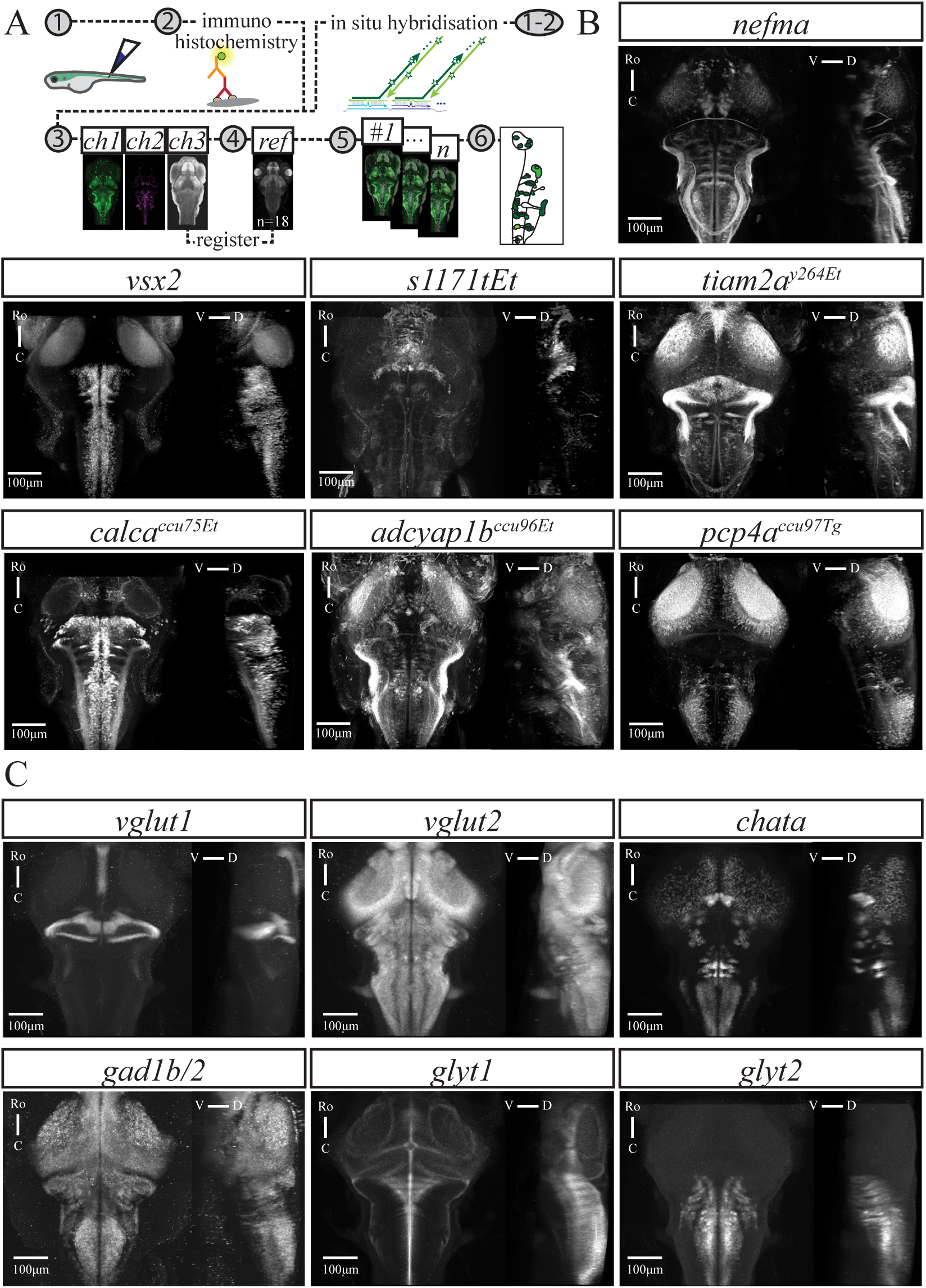
Overview of transgenic lines and neurotransmitter-associated gene expression patterns. A) Graphical summary of methods for reticulospinal backfills paired with immunohistochemistry (left) or *in situ* hybridisation (right), followed by confocal imaging, image registration to an average reference brain and analysis. B) GCaMP expression in transgenic lines used in this study as indicated in each panel. C) Expression of genes associated with neurotransmitter phenotypes as indicated in each panel. For B) and C) each panel shows a maximum intensity projection from the dorsal and sagittal view, taken from average stack following image registration. Number of fish: *nefma* n=7, *calca^ccu75Et^* n=12, *vsx2* n=11, *s1171tEt* n=5, *pcp4a^ccu97Tg^* n=6, *adcyap1b^ccu96Et^* n=8, *tiam2a^y264Et^* n=6, *vglut1* n=16, *vglut2* n=13, *chata* n=8, *gad1b/2* n=4, *glyt1* n=16, *glyt2* n=15. Scale bar 100*µ*m. For brain-wide expression patterns in example fish from selected transgenic lines see Fig. 1-1.

Here we provide an overview of the expression patterns of each transgenic line (Fig. 1-1). The *nefma* line has expression in the tectum, pretectum, tegmentum and hindbrain, as well as labelling the anterior and posterior lateral line ganglia, the trigeminal ganglion and neuromasts. Expression in the *vsx2* line is primarily in the hindbrain, whereas in the *s1171tEt* line expression is restricted to the tegmentum. The *tiam2a^y264Et^* line has expression in several areas throughout the brain, including the olfactory epithelium, optic chiasm, tectum, interpeduncular nucleus, tegmentum, hindbrain and cerebellum. The *calca^ccu75Et^* line mostly labels cells in the hindbrain, as well as a small number of cells in the mid- and forebrain, and outer retina. The *adcyap1b^ccu96Et^* line has expression in the olfactory epithelium, olfactory bulb, tegmentum, hindbrain, anterior and posterior lateral line ganglia, as well as sparse labelling in the tectum. Finally, the *pcp4a^ccu97Tg^* line has sparse labelling of cells in the forebrain and tectum, expression in the habenula, hindbrain, and the anterior and posterior lateral line ganglia.

For transgenic lines (*nefma, calca^ccu75Et^, s1171tEt*) that labelled cells beyond the classical RSNs we performed *in situ* hybridisation chain reaction (**isHCR**) to study the mRNA expression of different excitatory and inhibitory neurotransmitters: glutamate *(vglut1, vglut2a, vglut2b)*, acetyl choline transferase *(chata)*, gamma-aminobutyric acid *(GABA; gad1b, gad2)* and glycine *(glyt1, glyt2)* (Fig. 1B). The vesicular glutamate transporter *vglut2* has two orthologs in zebrafish, *vglut2a* and *vglut2b*, which had largely overlapping brain-wide expression of mRNA, consistent with data from the zebrafish brain atlas *mapzebrain* (Kunst et al., 2019). Except where explicitly stated, we refer to *vglut2* as a combination of *vglut2a* and *vglut2b*.

### The *nefma* line labels all RSNs and cranial nerve nuclei

We examined the organisation of RSNs in a knock-in reporter line in the *nefma* gene (Eschstruth et al., 2020) using dextran-conjugated retrograde labelling in larval zebrafish at 6 dpf (n=10). The *nefma* line has expression in the tectum, pretectum, tegmentum and hindbrain, as well as labelling the anterior and posterior lateral line ganglia, the trigeminal ganglion and neuromasts (Fig. 1-1). In this line, the Mauthner cell is not present at this age because it degenerates between 3-5 dpf (data not shown; Dr J. Bin and Dr D. Lyons, The University of Edinburgh, personal communication). The Mauthner homologues (MiD2cm, MiD2cl, MiD3cm, MiD3cl) remain intact. The back-labelled cells closely match the organisation previously described (Kimmel et al., 1982), which we term the ‘classical’ RSNs. All other classical RSNs that were back-labelled were present and expressing GCaMP (Fig. 2A-B). This includes the four named large cells (MeM1, MeLm, MeLr, MeLc) in the nucleus of the medial longitudinal fasciculus (**nMLF**), and RSNs in the hindbrain, including the rostral RoL1 and RoM1-3 cells, various medial cell groups (MiV1, MiV2, MiM1, MiR1, MiR2), and the caudal MiD2, MiD3 and MiT cells (Fig. 6-1). Note that the rostro-lateral RoL2-RoL3 and the caudal CaD and CaV cells were not successfully back-labelled, most likely requiring a different injection strategy. However, it is highly likely these cells are still present in the *nefma* line. We see numerous small GCaMP-positive cells lateral to the RoM3 cells, which could be the RoL2-3, and there are large GCaMP-positive cells situated caudally to the MiD3 cells most likely constituting the CaD/CaV cells (Kimmel et al., 1982; Orger et al., 2008).

**Figure 2:**
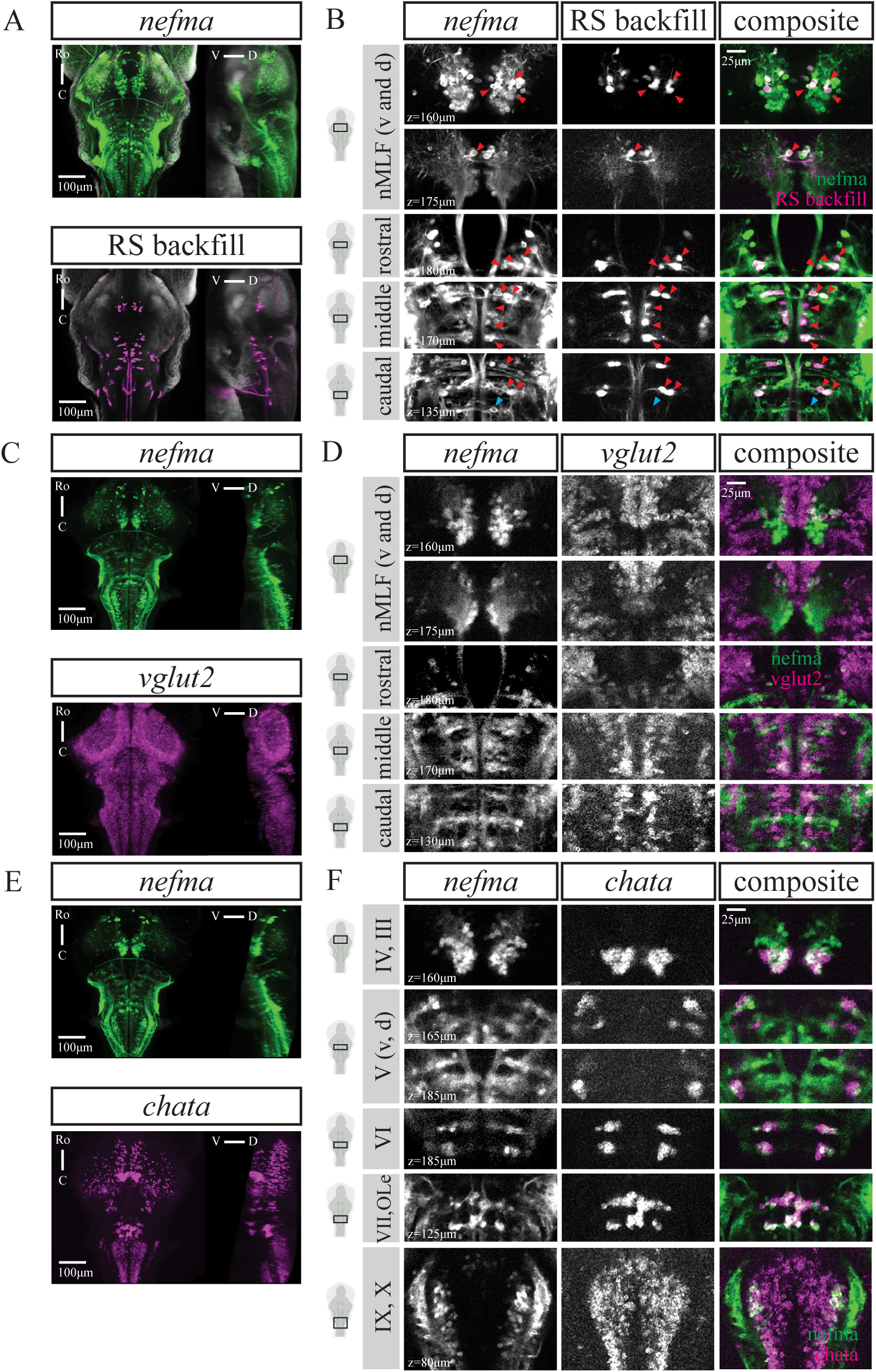
The *nefma* line labels all RSNs as well as cranial nerve nuclei. A) Maximum intensity projections (MIPs) from dorsal and sagittal view of an exemplary *nefma* fish (n=10) at 6 dpf with RSNs labelled via retrograde dye injection. B) Close ups at several planes to illustrate overlap between *nefma* line and backfill, indicated by red triangles. Putative CaD and CaV cells indicated by blue triangle. C) MIPs from dorsal and sagittal view of a *nefma* fish (n=24) at 6 dpf with *vglut2* mRNA expression. D) Close ups at several planes to illustrate considerable overlap between *nefma* line and *vglut2*. E) MIPs from dorsal and sagittal view of a *nefma* fish (n=16) at 6 dpf with *chata* mRNA expression. F) Close ups at several planes to illustrate cranial nerves III-VII, IX-X and octavolateralis efferent nucleus (OLe) are labelled in the *nefma* line. For A,C,E scale bar 100*µ*m, for B,D,F scale bar is 25*µ*m. For GABAergic (*gad1b/2*), glycinergic (*glyt1, glyt2*) or glutamatergic (*vglut1*) expression in neurons labelled by the *nefma* line see Fig. 2-1.

The majority of cells labelled by the *nefma* line, including the classical RSNs, are glutamatergic (*vglut2*; Fig. 2C-D) as confirmed by isHCR (n=24). In addition, the *nefma* line (n=16) labels cholinergic cells, which could be identified as the cranial nuclei (CNIII, oculomotor; CNIV, trochlear; CNV, trigeminal; CNVI, abducens; CNVII, facial; CNIX, glossopharyngeal; CNX, vagus) and octavolateralis efferent neurons (OLe; Fig. 2E-F), with locations consistent with anatomical annotations in zebrafish atlases (Kunst et al., 2019). There was no *vglut1*, GABAergic *(gad1b/2)* or glycinergic *(glyt1, glyt2)* mRNA expression in cells labelled by the *nefma* line (Fig. 2-1).

### Transgenic lines offer access to subpopulations of RSNs

#### calca^ccu75Et^

We established a new Gal4 transgenic line using a 5kb fragment of the *calca* promoter. The calca and adcyap1b promoters were cloned as part of a larger screen for short promoter sequences with neural expression, based on two characteristics: 1) annotation of expression in the hindbrain, among other brain regions (www.zfin.org, (Thisse et al., 2004)) and 2) a compact genomic locus (around 5kb), with a short first intron before the start codon that could be included within a short promoter sequence. The reason for this latter consideration is that many successful short promoter sequences used in zebrafish include the first intron, which has been associated with improved transgene expression (Higashijima et al., 1997). The *calca^ccu75Et^* line mostly labels cells in the hindbrain, as well as a small number of cells in the mid- and forebrain, and outer retina (Fig. 1-1). The organisation of RSNs in the *calca^ccu75Et^* line was examined using dextran-conjugated retrograde labelling at 6dpf (n=12). Many of the classical RSNs that were visualised by the dye injection into the rostral spinal cord were also labelled by the *calca^ccu75Et^* line (Fig. 3A-B). This includes the rostral RoM2, RoM3 and RoV3 cells, various medial cell groups (MiV1, MiV2, MiM1, MiR1, MiR2), and notably the caudal ipsilaterally-projecting (**IL**) MiD2i and MiD3i cells, but not their contralaterally-projecting (**CL**) counterparts. Following the projections of MiD2i and MiD3i at higher spatial resolution confirmed their identity as no decussation at the midline occurred (data not shown). In this line, the Mauthner cell is present but not labelled. For detailed RSN labelling across multiple fish see Fig. 6-3.

**Figure 3:**
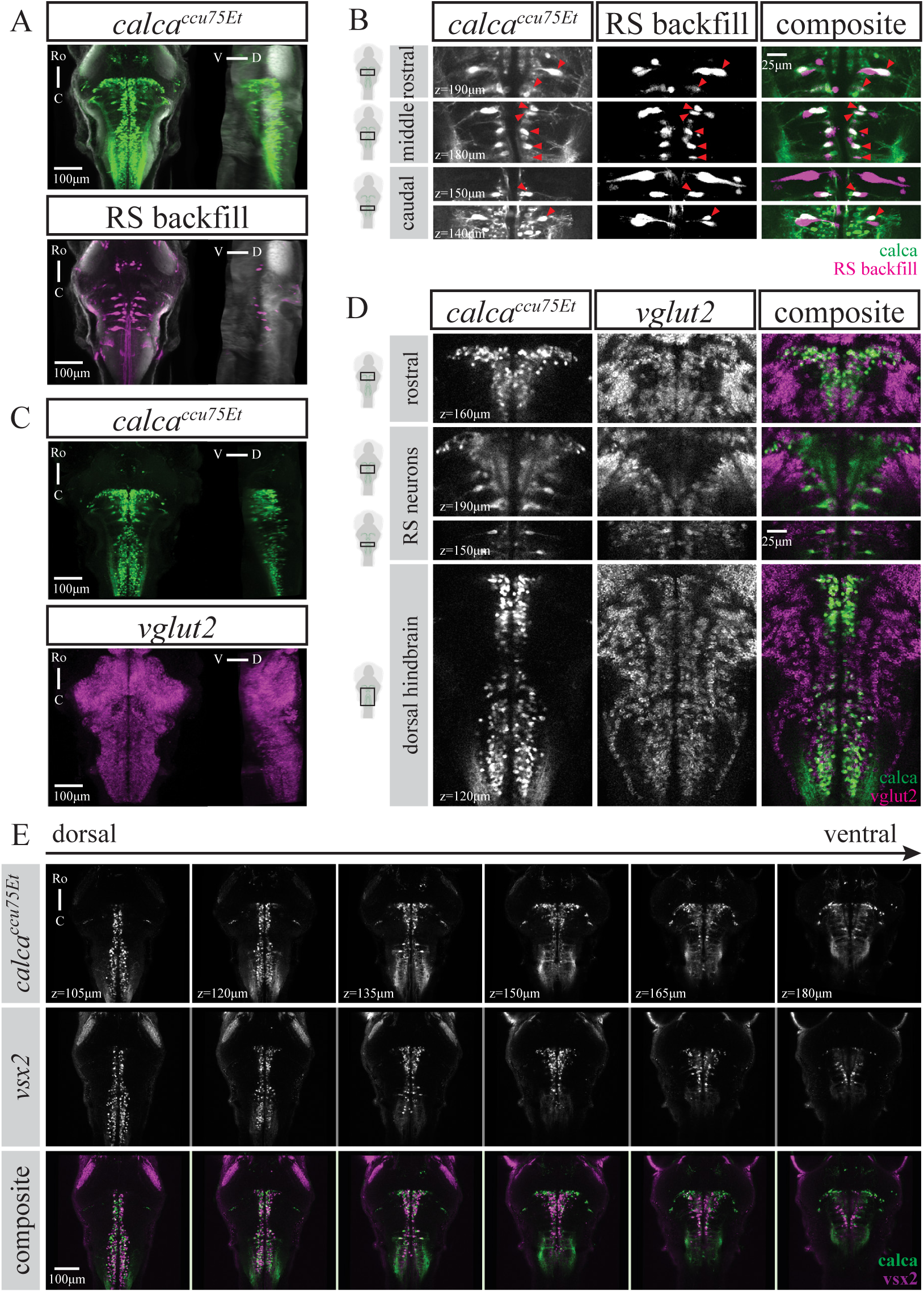
The *calca^ccu75Et^* line labels only IL-RSNs and closely matches the *vsx2* line. A) Maximum intensity projections from dorsal and sagittal view of an exemplary *calca^ccu75Et^* fish (of n=12) at 6 dpf with reticulospinal neurons labelled via retrograde dye injection. B) Close ups at several planes to illustrate overlap between *calca^ccu75Et^* line and backfill. Note that only IL-RSNs are present in *calca^ccu75Et^* as indicated by red triangles. C) Maximum intensity projections from dorsal and sagittal view of an exemplary *calca^ccu75Et^* fish (of n=8) at 6dpf with *vglut2* mRNA expression. D) Close ups at several planes to illustrate overlap between *calca^ccu75Et^* line and *vglut2*, particularly of the RSNs as well as a rostro-caudal glutamatergic (*vglut2*) stripe in the dorsal hindbrain. E) Single planes from dorsal (105*µ*m) to ventral (180*µ*m) at 15*µ*m intervals in *calca^ccu75Et^*, *vsx2* and composite. For both top and middle panel, images are from two single fish (of n=12 each) that were registered to a common reference brain (tERK). For A,C,E scale bar 100*µ*m, for B,D scale bar 25*µ*m. For cholinergic (*chata*), GABAergic (*gad1b/2*) or glycinergic (*glyt1, glyt2*) expression in neurons labelled by the *calca^ccu75Et^* line see Fig. 3-1.

There was no cholinergic (*chata*), GABAergic (*gad1b/2*) or glycinergic (*glyt1, glyt2*) mRNA expression in cells labelled by the *calca^ccu75Et^* line (Fig. 3-1). In fact, most cells labelled in the *calca^ccu75Et^* line were glutamatergic (*vglut2*), with a particularly prominent glutamatergic stripe spanning the whole medial dorsal hindbrain (Fig. 3C-D). This strongly resembled the well-characterised *vsx2* line that has a medial stripe of glutamatergic V2a neurons in the hindbrain (Kimura et al., 2006).

#### vsx2

We therefore compared the expression profiles of two *calca^ccu75Et^* and *vsx2* fish registered to a shared reference brain. This revealed a high degree of overlap between the two transgenic lines, with the exception of additional dorso-lateral transgene expression close to the midbrain boundary in the *calca^ccu75Et^* line (Fig. 3E-F).

To illustrate the similarity between the *calca^ccu75Et^* and *vsx2* lines at a cellular level, we also performed back-labelling of RSNs in *vsx2* fish at 6 dpf (n=12). Again, only ipsilaterally-projecting RSNs in the hindbrain were present in *vsx2* (Fig. 4A-B). This includes the rostral RoM2, RoM3 and RoV3 cells, various medial cell groups (MiV1, MiV2, MiM1, MiR1, MiR2), and the caudal ipsilaterally-projecting MiD2i and MiD3i cells. Following the projections of MiD2i and MiD3i at higher spatial resolution confirmed their identity with no decussation at the midline present (data not shown). Similarly to the *calca^ccu75Et^* line, the Mauthner cell was present but not labelled in *vsx2*. For detailed RSN labelling across multiple fish see Fig. 6-4.

**Figure 4:**
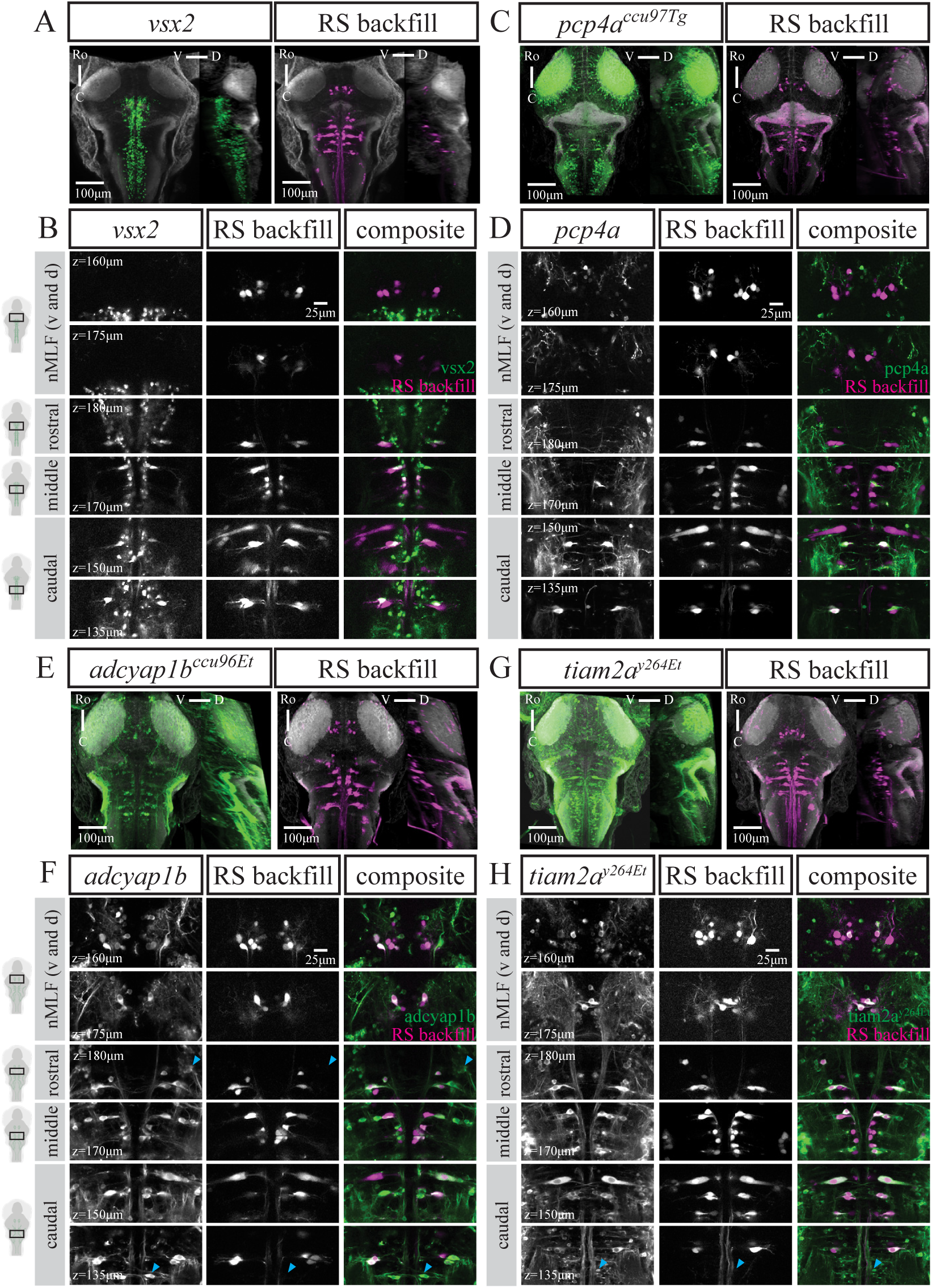
Reticulospinal cell labelling in four different transgenic lines. A,C,E,G) Maximum intensity projections from dorsal and sagittal view of an exemplary fish from A) *vsx2* (n=12), C) *pcp4a^ccu97Tg^* (n=12), E) *adcyap1b^ccu96Et^* (n=11) and G) *tiam2a^y264Et^* (n=12) at 6 dpf with RSNs labelled via retrograde dye injection. Scale bar is 100*µ*m. B,D,F,H) Close ups at several planes to illustrate overlap between B) *vsx2*, D) *pcp4a^ccu97Tg^*, F) *adcyap1b^ccu96Et^* or H) *tiam2a^y264Et^* and RS backfill. Blue triangle in F, rostral panel indicates putative RoL1. Blue triangle in F and H, caudal panel indicates putative CaD and CaV cells. Scale bar is 25*µ*m.

#### pcp4a^ccu97Tg^

We established a new transgenic line driving GAL4FF expression under the *pcp4a* promoter using a recombineered BAC. *Pcp4a* had been shown to be expressed in the telencephalon, habenula, pretectum, pre-glomerular complex, mammillary bodies, optic tectum and a subset of reticulospinal neurons (Mione et al., 2006). The *pcp4a^ccu97Tg^* line has sparse labelling of cells in the forebrain and tectum, expression in the habenula, hindbrain, and the anterior and posterior lateral line ganglia (Fig. 1-1). The reticulospinal back-labelling (n=12) specifically and reliably includes the dorso-caudal contralaterally-projecting MiD2cm, MiD2cl, MiD3cm and MiD3cl but not their ipsilateral counterparts (Fig. 4C-D). Labelling of other RSNs in this line was more variable, with different combinations of various medial cell groups (MiV1, MiV2, MiM1, MiR1, MiR2) being labelled in 50% of fish (Fig. 6-6). To our knowledge, this is the first transgenic line specifically labelling the contralaterally-projecting MiD cells of the Mauthner array.

#### adcyap1b^ccu96Et^

We established a new transgenic line in which GAL4FF expression is driven by a 1.7kb promoter region upstream of the start codon of adenylate cyclase activating polypeptide 1b (*adcyap1b*) gene. The *adcyap1b* gene encodes pituitary adenylate cyclase-activating polypeptide 2 (**PACAP2**), is evolutionarily conserved and expressed in the telencephalon, diencephalon, rhombencephalon, and dorsal spinal cord (Alexandre et al., 2011). PACAP2 plays a role in brain development (Wu et al., 2006) and the neuropeptide *adcyap1b* has been shown to enhance sensory responsiveness in larval zebrafish (Woods et al., 2014). *adcyap1b* was included in the previously mentioned screen based on the same characteristics described above for calca. In addition to labelling cells in the tegmentum and hindbrain, our *adcyap1b^ccu96Et^* line has expression in the olfactory epithelium, olfactory bulb, anterior and posterior lateral line ganglia, as well as sparse labelling in the tectum (Fig. 1-1). Following back-filling with dextran-conjugated tracer dye (n=11), we report overlap with GCaMP-expression in the four identified cells of the nMLF (MeM1, MeLm, MeLr, MeLc), the Mauthner cell and most other RSNs including the vestibular cells (Fig. 4E-F). We observed labelling of most of the caudal contralaterally-projecting MiD cells (MiD2cm, MiD2cl, MiD2i, MiD3cm and MiD3cl) but not the MiD3i and MiT cells. For detailed RSN labelling across multiple fish see Fig. 6-5. As mentioned previously, the CaD, CaV, RoL1 and RoL2 cells were not retrogradely labelled in our preparation. However, it is likely that the *adcyap1b^ccu96Et^* line labels those cells well, as there are large cell bodies visible in their respective putative locations (Kimmel et al., 1982; Orger et al., 2008).

#### tiam2a^y264Et^

The previously established *tiam2a^y264Et^* line (Marquart et al., 2015) was of interest due to labelling the Mauthner cell and homologues. It has expression throughout the brain, including the olfactory epithelium, optic chiasm, tectum, interpeduncular nucleus, tegmentum, hindbrain and cerebellum **(Fig. 1-1)**. Backfills (n=12) revealed that this line labelled MeM1 more reliably than the other identified nMLF neurons (MeLm, MeLr, MeLc), as well as consistently labelling the rostral RoM2l, RoM3m and RoM3l, the Mauthner cell, and both the CL- and IL-projecting MiD2 and MiD3 cells in the hindbrain (Fig. 4G-H). Labelling of other RSNs was more variable across fish (Fig. 6-7). Similarly to above, it is likely the CaD and CaV cells are also labelled in the *tiam2a^y264Et^* line (Kimmel et al., 1982; Orger et al., 2008).

### The *s1171tEt* line labels *vglut2*-expressing neurons in the nMLF

The organisation of midbrain RSNs in the *s1171tEt* line was examined using dextran-conjugated retrograde labelling in larval zebrafish at 6dpf (n=12). The *s1171tEt* has a triangular expression profile centred around the nMLF in the tegmentum, rostral to the cholinergic oculomotor nucleus (Thiele et al., 2014). It includes the four canonical large nMLF cells (MeM1, MeLm, MeLr, MeLc), as well as small mesencephalic cells (**MeS**; Fig. 5A-B). For detailed RSN labelling across multiple fish see Fig. 6-2.

**Figure 5:**
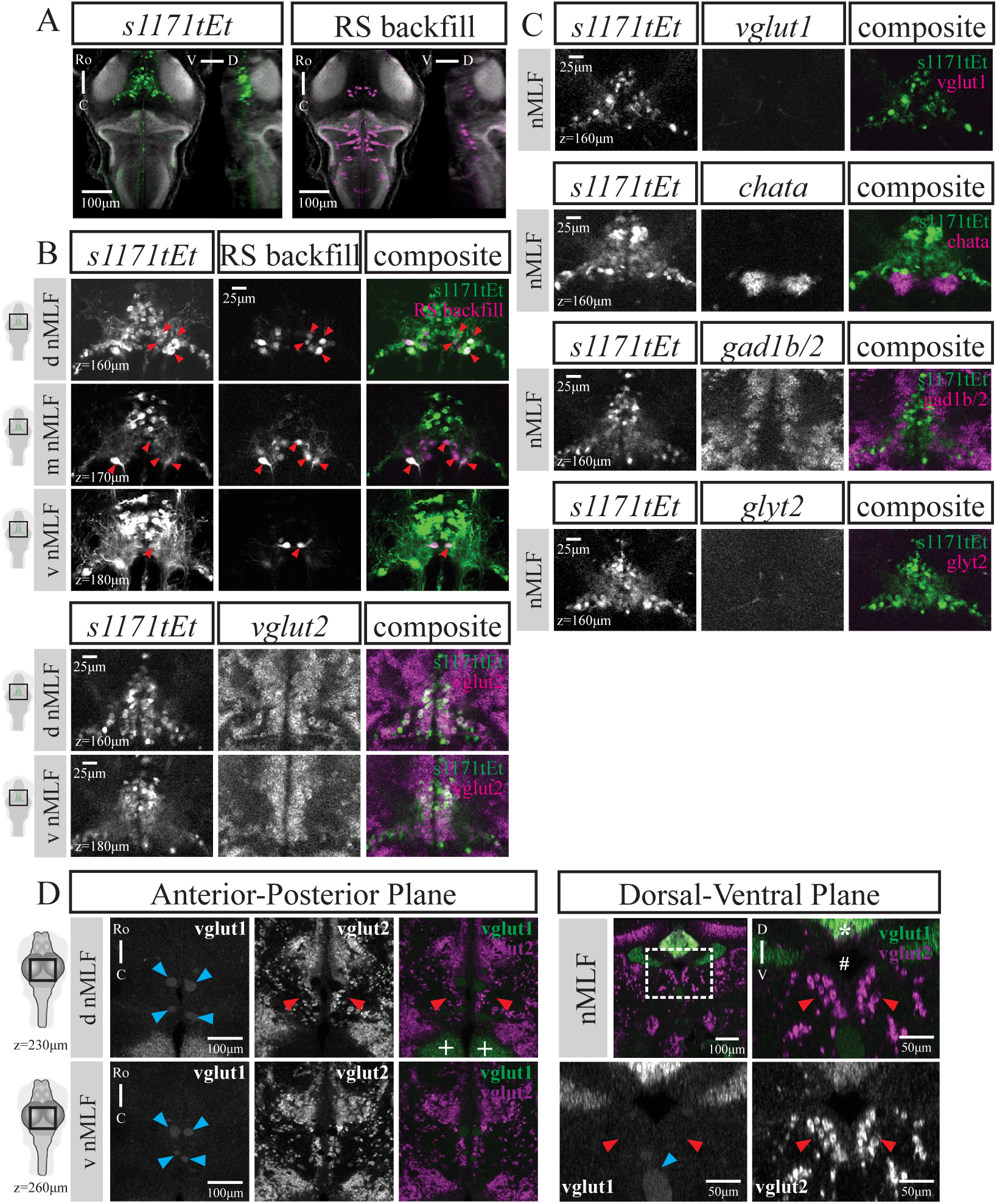
The *s1171tEt* line labels RSNs in the nMLF and is glutamatergic (*vglut2*) across development. A) Dorsal and sagittal maximum intensity projections from an example *s1171tEt* fish (of n=12) at 6 dpf with RSNs labelled via retrograde dye injection. Scale bar 100*µ*m. B) Close ups at several planes to illustrate overlap between *s1171tEt* line and backfill, note the four main RSNs as indicated by red triangles. Scale bar 25*µ*m. C) Close ups at several planes of exemplary fish to show considerable overlap between *s1171tEt* and *vglut2* (n=8), particularly in RSNs. No overlap between *s1171tEt* and *vglut1* (n=7), *chata* (n=8), *gad1b/2* (n=8) or *glyt2* (n=7). Scale bar 25*µ*m. D) *vglut1* and *vglut2* expression in exemplary 4 week-old juvenile fish (n=4). Left) two planes in dorsal view, scale bar 100*µ*m. Right) matching transverse view, scale bar 50*µ*m. Blue triangles indicate blood vessels, red triangles point towards region of nMLF. For more examples of glutamatergic expression patterns in 4 week-old juvenile fish see Fig. 5-1.

The majority of cells in the *s1171tEt* line were glutamatergic, specific for the vesicular glutamate transporter 2 (*vglut2*; Fig. 5B). We did not detect *vglut1*-expressing cells near the nMLF, as well as no cholinergic (*chata*), GABAergic (*gad1b/2*) or glycinergic (*glyt1, glyt2*) mRNA expression in cells labelled by the *s1171tEt* line (Fig. 5C). A previous study distinguished lateral and medial regions of the nMLF in zebrafish at 4-weeks post-fertilisation based on different levels of expression observed in *vglut1* and *vglut2a* transgenic lines (Berg et al., 2023). This difference with the isHCR results could reflect a change in the expression pattern over development, or alternatively could be due to expression in these transgenic lines that does not reflect the natural gene expression patterns.

To understand this, we asked whether glutamatergic expression profiles change across development. We performed a modified *in situ* hybridisation chain reaction protocol on brains dissected from juvenile zebrafish (n=4; 4-weeks post-fertilisation). Using landmarks identified with the adult zebrafish brain atlas (**AZBA** (Kenney et al., 2021)) —such as the torus longitudinalis (**TL**), valvula cerebelli, and the diencephalic ventricle —allowed us to identify several brain regions. Glutamatergic expression appears stable across development, with *vglut1* expression being restricted to the TL and the cerebellum, in contrast to brain-wide *vglut2* expression, including the putative nMLF (Fig. 5-1). To identify the nMLF in the juvenile brain, we again utilised the AZBA. Matching the triangular shape of the nMLF on AZBA in the transverse view, situated between the diencephalic ventricle and a cluster of *vglut2* cells most likely belonging to the red nucleus, is a cluster of large *vglut2* cells characteristic of the nMLF (Fig. 5D). We did not observe *vglut1* expression near this cluster of cells, only auto-fluorescent blood vessels.

### Summary of RSN labelling across transgenic lines

The transgenic lines characterised in this study can be utilised in a complementary fashion to study reticulospinal circuits at larval stages (Fig. 6). The *nefma* line (n=10) reliably labels all RSNs. However, this line cannot be used to study the Mauthner cell since it degenerates early in development. The *adcyap1b^ccu96Et^* line (n=11) labels most RSNs including the Mauthner cell. However, the MiD3i and MiT cells are not labelled, and we emphasise the need for rigorous pre-screening due to silencing. The most suitable line to study Mauthner and homologue activity is the *tiam2a^y264Et^* line (n=12), with labelling of other RSNs being more variable. The *calca^ccu75Et^/vsx2* and *pcp4a^ccu97Tg^* lines offer an interesting opportunity by giving genetic access to complementary projection patterns in the Mauthner array, with the *calca^ccu75Et^* line (n=12) only labelling the ipsilaterally-projecting MiD cells (as well as other rostral and medial RSNs) and the *pcp4a^ccu97Tg^* line (n=12) specifically labelling contralaterally-projecting MiD cells, while labelling of various medial RSNS is much more rare. In addition, we have demonstrated that the *calca^ccu75Et^* line closely matches the existing *vsx2* line (n=12), both in broad expression pattern and at the single-RSN-level – though it labels additional cells in the rostrolateral hindbrain. For exact numbers across fish for all transgenic lines, see Extended Data 6-1 until 6-7). Finally, for wider nMLF-related studies the *s1171tEt* line (n=12) offers broad labelling in the tegmentum, including the four canonical nMLF-RSNs (MeM1, MeLr, MeLc, MeLm).

**Figure 6:**
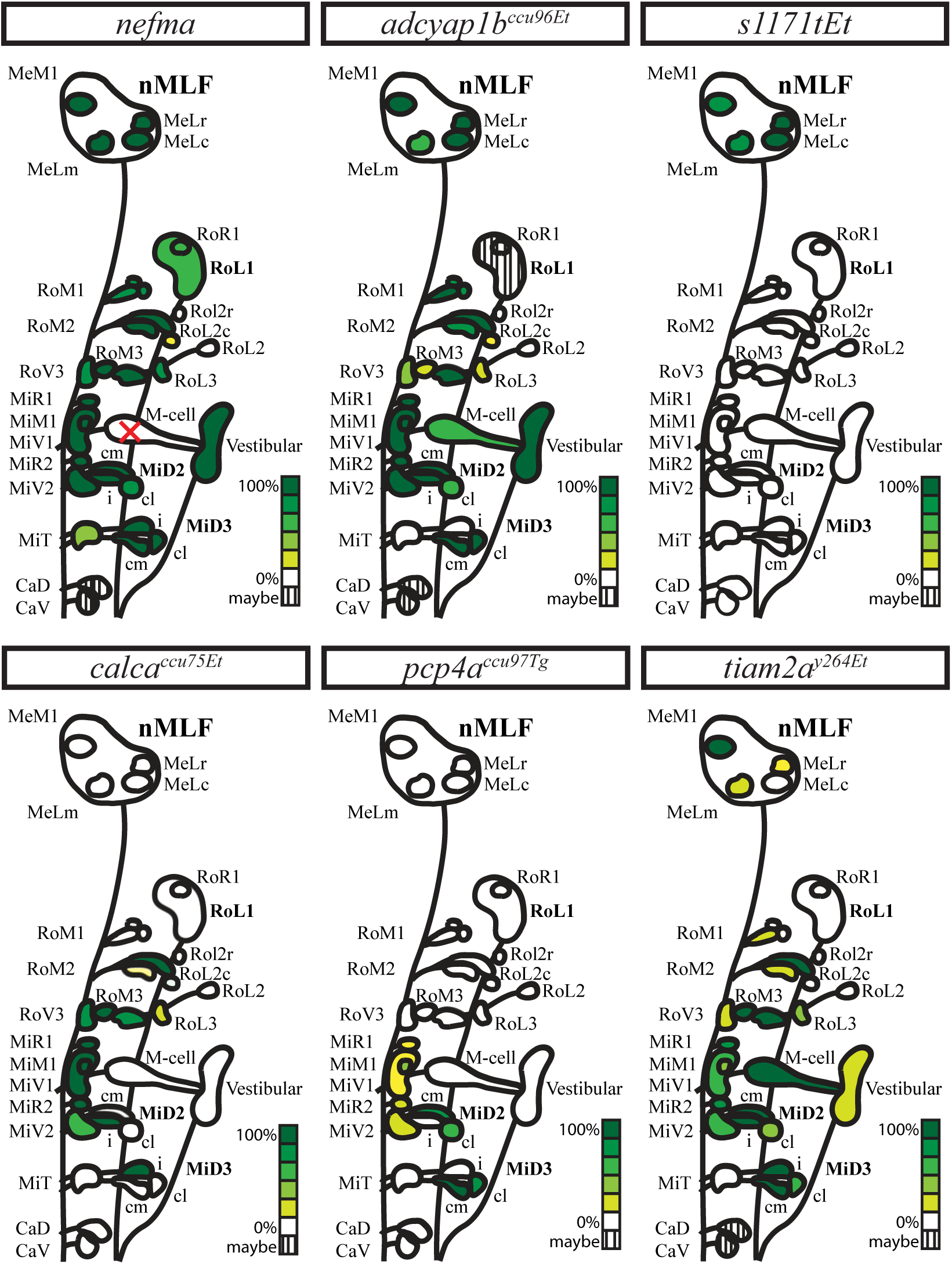
Graphical summary of the number of RSNs labelled in each transgenic line across multiple fish. *nefma* n=10, *calca^ccu75Et^* n=12, *vsx2* n=12, *s1171tEt* n=12, *pcp4a^ccu97Tg^* n=12, *adcyap1b^ccu96Et^* n=11, *tiam2a^y264Et^* n=12. For data on which reticulospinal cells are labelled in each fish per transgenic line see Fig. 6-1 until Fig. 6-7.

We further demonstrate that cells labelled in these transgenic lines are glutamatergic (*vglut2*), with the *nefma* line additionally labelling the cholinergic cranial neurons.

## Discussion

The aim of this study was to provide a detailed account of seven transgenic lines with gene expression in the brainstem of larval zebrafish. We characterized four existing transgenic lines *(nefma, vsx2, s1171tEt, tiam2a^y264Et^*) and presented three newly established transgenic lines *(calca^ccu75Et^, pcp4a^ccu97Tg^, adcyap1b^ccu96Et^)*. For each transgenic line, we performed retrograde labelling of reticulospinal neurons (**RSNs**) followed by immunostaining against GCaMP/RFP and the whole-brain neural marker total ERK for subsequent image registration purposes.

As previously reported (Thiele et al., 2014) the *s1171tEt* line reliably labels the four large cells (MeLr, MeLc, MeLm, MeM1) in the nucleus of the medial longitudinal fasciculus (**nMLF**) as well as many other neurons in the surrounding area. We observed occasional overlap with other backfilled RSNs, most likely constituting the small mesencephalic cells (**MeS**), first characterized by (Thiele et al., 2014). The nMLF in fish is thought to be homologous to the interstitial nucleus of Cajal (Yamamoto et al., 2017; Carbo-Tano et al., 2023), or a part of such a structure (Wullimann et al., 1996), due to its involvement in postural control and projections to the spinal cord during early development (Thiele et al., 2014; Sugioka et al., 2023; Liu and Bagnall, 2023).

Two transgenic lines labelled almost all RSNs, with notable exceptions: the *nefma* line reliably labels all RSNs except for the Mauthner cell, which degenerates earlier in development. We can speculate as to why the Mauthner cell degenerates in this transgenic line. The degeneration appears to be independent of the presence of a UAS reporter, consistent with reports that the KaLTA4 construct (Distel et al., 2009), can be cytotoxic at high concentrations (Zhang et al., 2019). An alternative explanation is that high axon calibre neurons like the Mauthner cell are more sensitive to perturbations of *nefma* gene expression that arise from the insertion of the transgene in that locus (Baraban et al., 2013). Another transgenic line has been made by insertion into the same locus (Liu et al., 2020), in which the Mauthner cell does not degenerate (Hamling et al., 2023). Multiple differences exist between the two lines, which could account for the discrepancy. First, in the line used here, KalTA4 is integrated in frame at the end of the *nefma* gene with a self-cleaving peptide sequence between the two, while in the other line the construct is integrated upstream of the *nefma* gene. Second, a different GAL4 construct was used, with the other line using GAL4FF rather than KalTA4. It is likely that the other line has a similar expression pattern, with most RSNs labelled, but this remains to be systematically documented. We were also not able to confirm labelling of the rostral RoL2-3 and caudal CaD and CaV neurons, as these were not successfully labelled by backfilling in our preparation and may require a different injection strategy. However, there are GCaMP-expressing cells in the *nefma* line where we would expect the RoL2-3 and CaD and CaV cells to be based on previous studies (Kimmel et al., 1982; Orger et al., 2008). We thus conclude that the *nefma* line labels all RSNs except the Mauthner cell. The newly generated *adcyap1b^ccu96Et^* line reliably labels most RSNs including the Mauthner cells and most homologues and the putative CaD and CaV cells, with a specific exception: we did not observe labelling of the MiT and ipsilaterally-projecting MiD3i cells in the *adcyap1b^ccu96Et^* line.

To study the Mauthner array, three complementary transgenic lines can be used. The *tiam2a^y264Et^* line reliably labels the Mauthner cells and homologues (including the MiD2i and MiD3i), however, labelling of other RSNs is stochastic, as previously reported by (Tabor et al., 2014). We presented two new lines that offer complementary genetic access to different projection patterns within the Mauthner array. The *calca^ccu75Et^* line largely overlaps with the well-known line *vsx2*, with the addition of gene expression in the rostral lateral hindbrain. The *calca^ccu75Et^* line reliably labels the RoM2, RoM3 and various medial RSNs as well as the ipsilaterally-projecting Mauthner homologues MiD2i and MiD3i. Conversely, the *pcp4a^ccu97Tg^* line reliably labels the contralaterally-projecting medial and lateral MiD2cm, MiD2cl, MiD3cm, MiD3cl as well as more rarely various medial cells. To our knowledge, this is the first description of a transgenic line in zebrafish providing specific genetic access to the contralaterally-projecting Mauthner homologues.

There is ample evidence for the usefulness of transgenic lines labelling different subpopulations in the hindbrain to dissect premotor circuitry and function. For instance, using a Gal4-transgenic line under the *nefma* promoter allowed experimenters to reliably record convergent afferent signals in vestibulospinal neurons in vivo (Liu et al., 2020). It was later shown in the same line that vestibulospinal neuron activity increases as a function of ipsilateral tilt amplitude using tilt-in-place microscopy (Hamling et al., 2023).

Another transgenic line identified in a Gal4 enhancer trap screen, *s1171tEt*, with expression in the thalamus, cerebellum and trunk musculature (Scott and Baier, 2009) was instrumental in showing that the four large (MeLr, MeLc, MeLm, MeM1) as well as smaller (MeS) nMLF neurons are involved in steering movements (Thiele et al., 2014), tail bending (Dal Maschio et al., 2017) and relay commands from the pretectal AF7 to the hindbrain, potentially involved in hunting behaviour (Semmelhack et al., 2014). In addition to their terminal arbours in the spinal cord, it was also demonstrated that nMLF cells labelled in the *s1171tEt* line have extensive axon collaterals in the hindbrain, suggesting they may be in involved in coordination of different premotor regions (Thiele et al., 2014).

Information on RSNs involved in steering behaviours also comes from another study using a transgenic line that labels V2a neurons, showing they receive innervation from the mesencephalic locomotor region (**MLR**) and are also involved in forward swimming (Carbo-Tano et al., 2023). Initially generated as a BAC line named chx10 after the mammalian homologue, it has been named *alx* and *vsx2* in fish. It labels medial ipsilaterally-projecting cells in the hind-brain and spinal cord, shown to be glutamatergic descending interneurons providing excitatory drive to motoneurons in the spinal cord (Kimura et al., 2006). It was subsequently confirmed that they overlap with the most medial V2a glutamatergic stripe in the hindbrain (Kinkhabwala et al., 2011). Optogenetic activation of *chx10*-expressing neurons in the hindbrain using channelrhodopsin evoked swimming, while forced inactivation using Archearhodopsin3 or Halorhodopsin reliably stopped ongoing swimming (Kimura et al., 2013).

Finally, the *tiam2a^y264Et^* line labels the Mauthner array reliably, and other RSNs more stochastically, as well as the anterior lateral line ganglia amongst others. Studies using the *tiam2a^y264Et^* line showed that direct activation of the Mauthner cell by electric field pulses could drive ultrarapid escape responses (Tabor et al., 2014), and that it receives pre-pulse inhibition from gsx2-glutamatergic neurons (Tabor et al., 2018).

Together, these studies illustrate the usefulness of transgenic lines labelling different sets of reticulospinal neurons in order to decipher their role in movement control. Another option to record neural activity in this population is to retrogradely label the neurons by spinal injections of a dextran conjugated fluorescent calcium indicator dye, such as Cal520 (Tada et al., 2014). However, spinal injections can be variable, require experimental skill and are time consuming. Importantly, their invasive nature, which necessarily damages the axons of the cells of interest, has the potential to impact the fish’s behaviour, altering the circuits that we wish to study (Gahtan and O’Malley, 2001). To complement functional imaging studies, the transgenic lines described here can be used to express optogenetic tools to perturb neuronal function, enabling optical manipulation of cell activity. Since all of the lines have expression in multiple targets, some additional spatial targeting of the incident light will be needed to achieve specificity. This has been achieved in practice in zebrafish through the use of optical fibres (Arrenberg et al., 2009), patterned projections (Zhu et al., 2012; Wyart et al., 2009) and 2-photon holography (Dal Maschio et al., 2017). Despite this caveat, restriction of expression to a small subset of possible neurons will minimize off-target effects and simplify experimental design. Other tools can benefit from similar intersections of optical and genetic specificity. For example, the photosensitizer KillerRed can be used to target cells for light-mediated ablation (Teh et al., 2010), while optical highlighter proteins can be used to reveal single neuron morphologies and projections with minimal background (Datta and Patterson, 2012).

In addition to reticulospinal backfills, we confirmed that cells labelled by the *nefma*, *calca^ccu75Et^* and *s1171tEt* transgenic lines are glutamatergic (*vglut2*). The *nefma* line also labels the cholinergic cranial motor neurons III-VII and IX-X. A previous study had reported medial *vglut2*-positive and lateral *vglut1*-positive cells in the nMLF in zebrafish at 4-6 weeks-post-fertilisation based on expression patterns of transgenic lines (Berg et al., 2023). We did not observe *vglut1*-positive cells in the nMLF in our study on larval zebrafish, instead, *vglut1*-expression was restricted to the torus longitudinalis (**TL**) and cerebellum. This finding is corroborated by isHCR data available in the larval zebrafish atlas *mapzebrain* (Kunst et al., 2019). We thus wanted to understand whether glutamatergic expression profiles change over development. We performed isHCR against *vglut1* and *vglut2* in brains dissected from 4 week old zebrafish, an age consistent with the study by Berg et al. (Berg et al., 2023). Similarly to the larvae, we only observed *vglut1*-expression in the TL and cerebellum, identified using the adult zebrafish brain atlas *AZBA* (Kenney et al., 2021). *Vglut2*-expression was more widespread, including expression in a region likely constituting the nMLF – identified using landmarks, such as the diencephalic ventricle, from AZBA. Again, we wondered whether glutamatergic expression profiles in the nMLF would change at later stages of development. However, this does not seem to be the case: a study in adult zebrafish (older than 90 days) using *in situ* hybridisation reports *vglut1*-expression in the TL and cerebellum only (Bae et al., 2009). We therefore think that the distinction between nMLF subregions in Berg *et al*. (2023) reflects variation in transgenic expression patterns, rather than the natural expression of these genes. However, this hypothesis needs to be tested by directly comparing *in situ* hybridisation data with expression in those transgenic lines.

In conclusion, we have provided a detailed account of the degree of RSN labelling in seven transgenic lines with glutamatergic (*vglut2*) expression in the brainstem in larval zebrafish, offering projection-specific genetic access to subpopulations of RSNs. This resource provides a useful basis for future research, including the possibility to combine with genetically-encoded activity reporters and optogenetic tools, in the hope to uncover fundamental principles in the descending control of locomotion.

## Author Contributions

EC and MO designed research; EC, PS and AO performed research; SR, ART and RD contributed unpublished reagents/analytic tools; EC and AO analyzed data; EC and MO wrote the paper. All authors read and approved the manuscript.

## Acknowledgements

We thank Dr. S. Lackner (Champalimaud Foundation) for cloning of the calca promoter fragment for Tg[-5.0calca:Gal4FF^ccu75Et^]. We thank I. Oliveira and J. Monteiro (Champalimaud Foundation) for preliminary imaging of the Tg[nefma:KalTa4] line. We thank the Champalimaud Foundation Advanced Bio-imaging and Bio-optics Experimental platform (ABBE) for technical support with confocal imaging and the Champalimaud Foundation Fish Platform for meticulous fish care. We thank Dr Chanpreet Singh (Molecular Instruments) for technical advice on adapting the *in situ* hybridisation protocol to juvenile fish. We thank J. Bin and D. Lyons (University of Edinburgh, UK) for sharing unpublished data on the Mauthner cell degeneration in Tg[nefma:KalTa4]. We would like to thank the Giudicelli lab (Sorbonne University, France) for generating and sharing embryos from Tg[nefma:KalTa4], the Schuster lab (University of Bayreuth, Germany) for sending embryos from Tg[tiam2a^y264Et(B)^] and the Wyart lab (Paris Brain Institute, France) for sending embryos from TgBAC[vsx2:Gal4FF^nns18Tg^, UAS:mRFP].

## Extended Data

**Figure 1-1:**
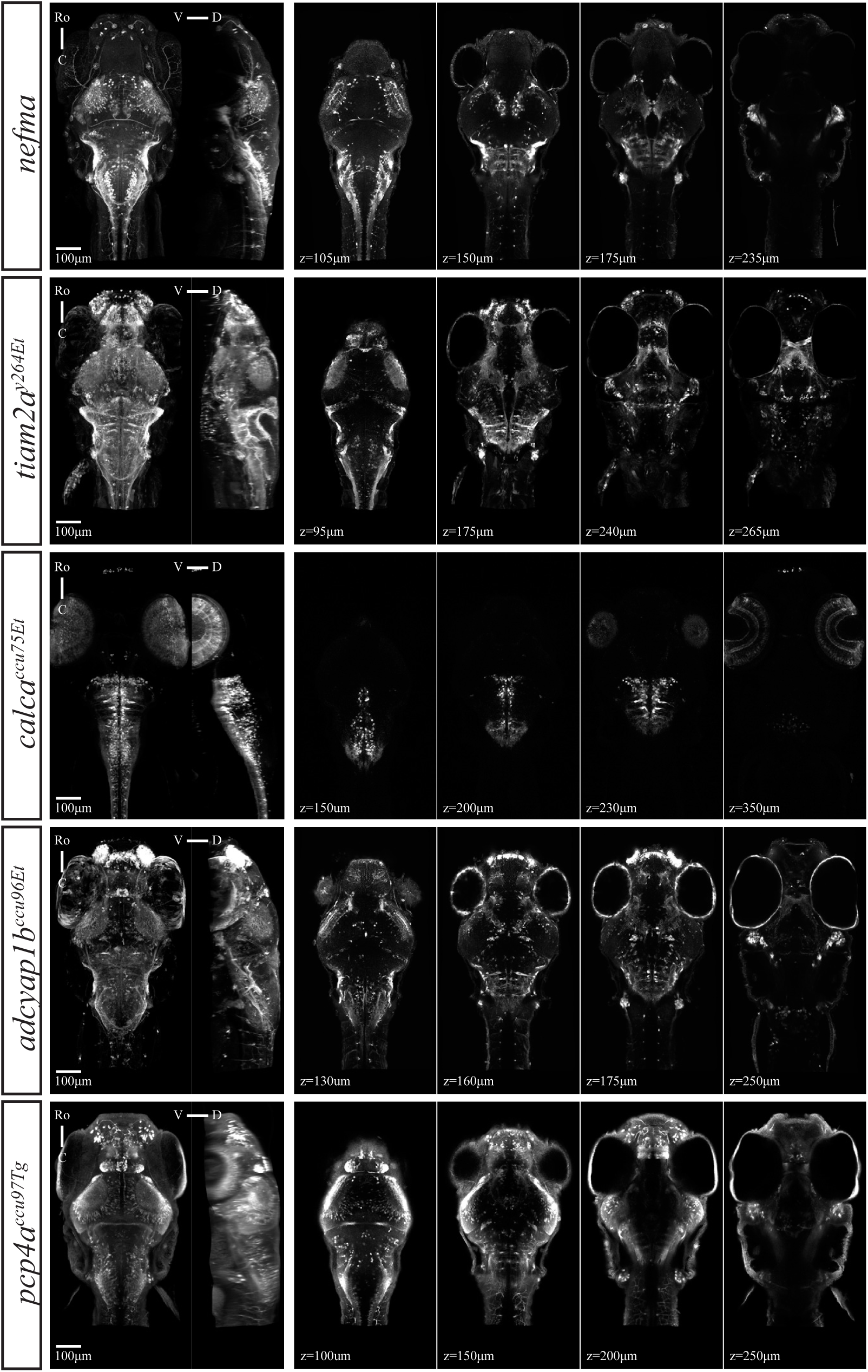
Example fish to show brain-wide expression patterns in selected transgenic lines. The *nefma* line has expression in the tectum, pre-tectum, tegmentum and hindbrain, as well as labelling the anterior and posterior lateral line ganglia, the trigeminal ganglion and neuromasts. The *tiam2a^y264Et^* line has expression in the olfactory epithelium, optic chiasm, tectum, interpeduncular nucleus, tegmentum, hindbrain and cerebellum. The *calca^ccu75Et^* line mostly labels cells in the hindbrain, as well as a small number of cells in the mid- and forebrain, and outer retina. The *adcyap1b^ccu96Et^* line has expression in the olfactory epithelium, olfactory bulb, tegmentum, hindbrain, anterior and posterior lateral line ganglia, as well as sparse labelling in the tectum. The *pcp4a^ccu97Tg^* line has sparse labelling of cells in the forebrain and tectum, expression in the habenula, hindbrain, and the anterior and posterior lateral line ganglia. Scale bar is 100*µ*m.

**Figure 2-1:**
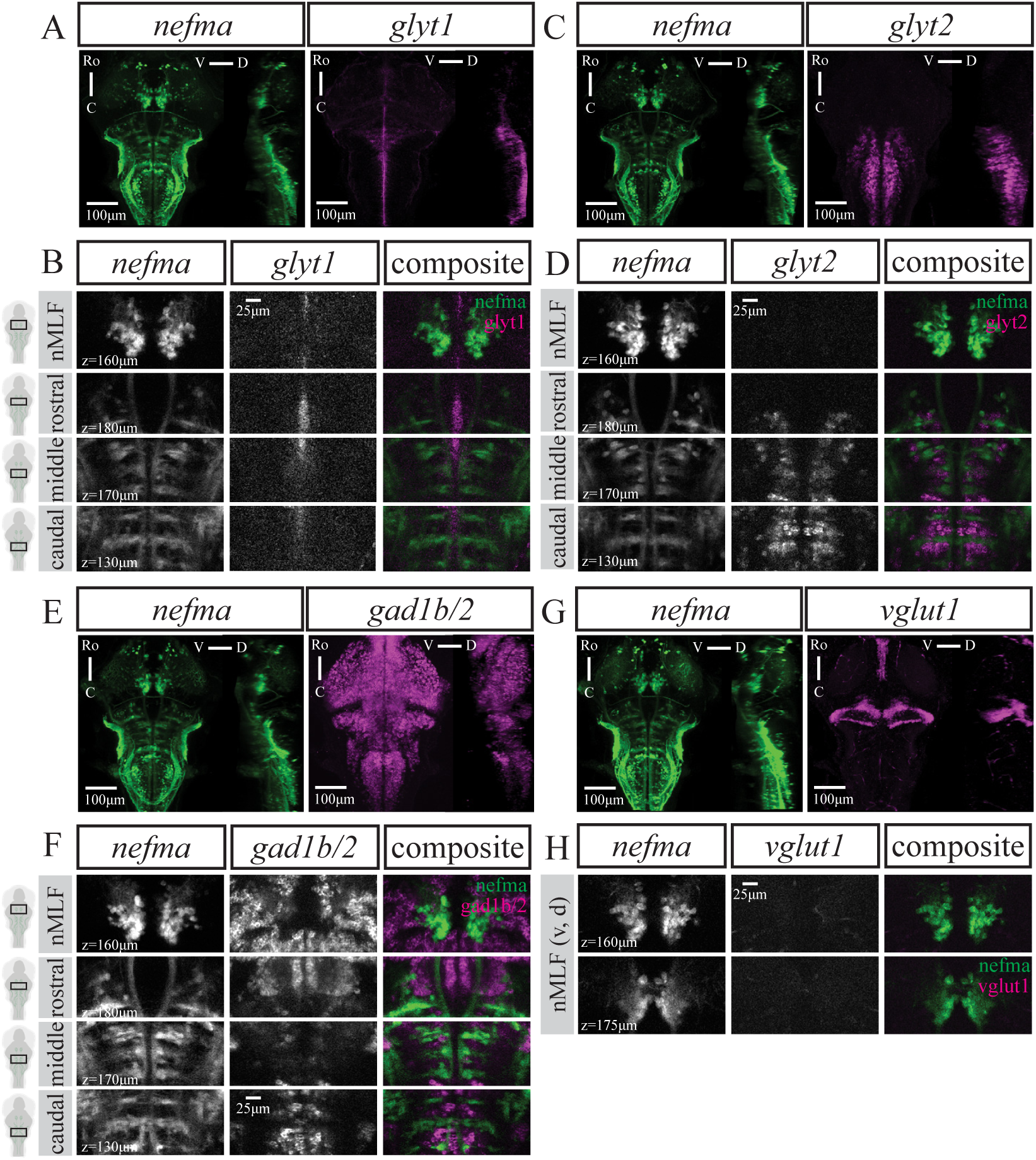
No GABAergic (*gad1b/2*), glycinergic (*glyt1, glyt2*) or glutamatergic (*vglut1*) expression in neurons labelled by the *nefma* line. A,C,E,G) Maximum intensity projections from dorsal and sagittal view of an exemplary *nefma* fish at 6dpf with A) *gad1b/2* (n=8), C) *glyt1* (n=16), E) *glyt2* (n=15) or G) *vglut1* (n=14) mRNA expression. Scale bar is 100*µ*m. Close ups at several planes to better illustrate no overlap between *nefma* line and B) *gad1b/2*, D) *glyt1*, F) *glyt2* or H) *vglut1* mRNA-expressing neurons. Scale bar is 25*µ*m.

**Figure 3-1:**
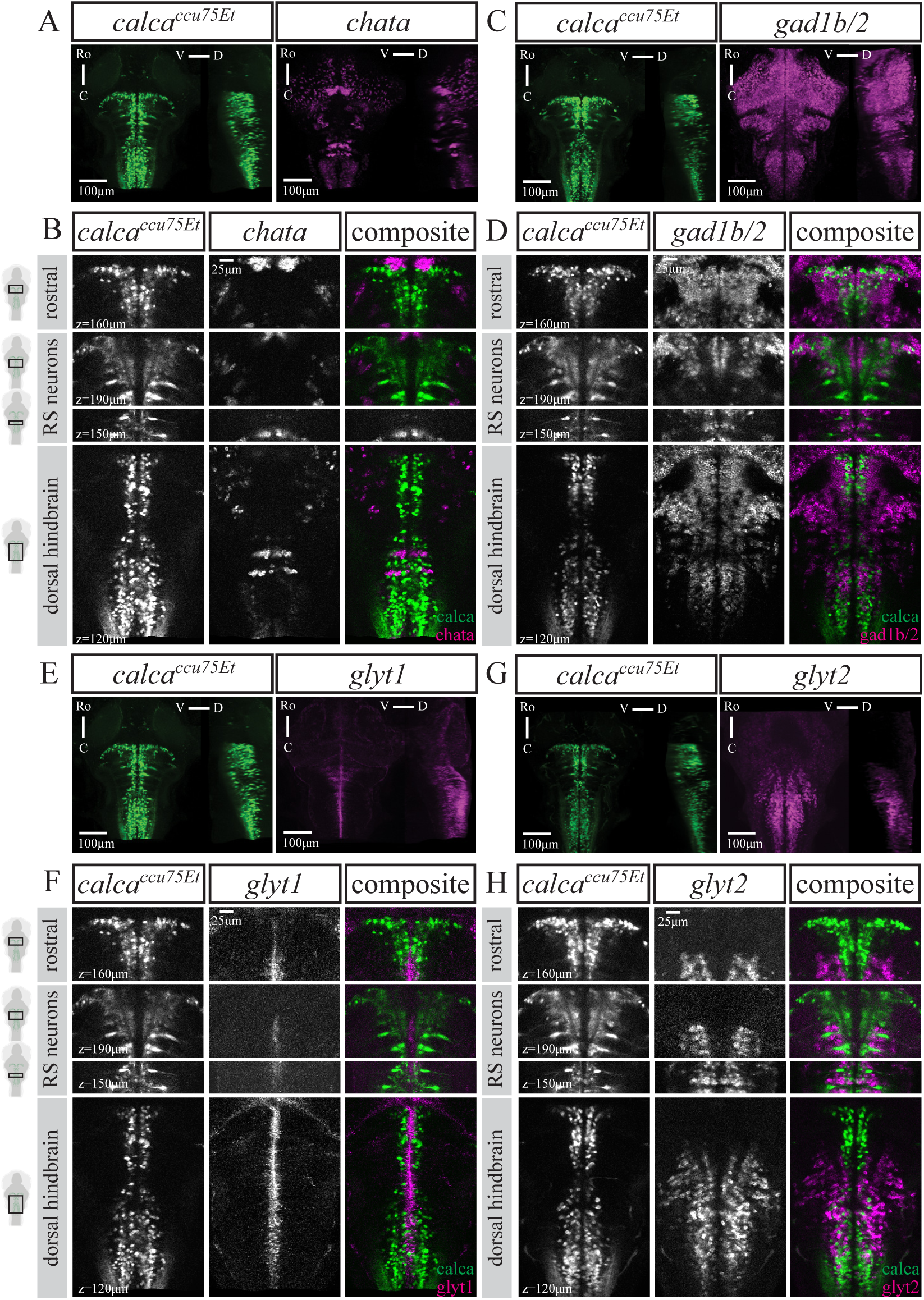
No cholinergic (*chata*), GABAergic (*gad1b/2*) or glycinergic (*glyt1, glyt2*) expression in neurons labelled by the *calca^ccu75Et^* line. A,C,E,G) Maximum intensity projections from dorsal and sagittal view of an exemplary *calca^ccu75Et^* fish at 6dpf with A) *chata* (n=15), C) *gad1b/2* (n=7), E) *glyt1* (n=16), G) *glyt2* (n=16) mRNA expression. Scale bar is 100*µ*m. B,D,F,H) Close ups at several planes to better illustrate no overlap between *calca^ccu75Et^* line and B) *chata*, D) *gad1b/2*, F) *glyt1* or H) *glyt2* or mRNA-expressing neurons. Scale bar is 25*µ*m.

**Figure 5-1:**
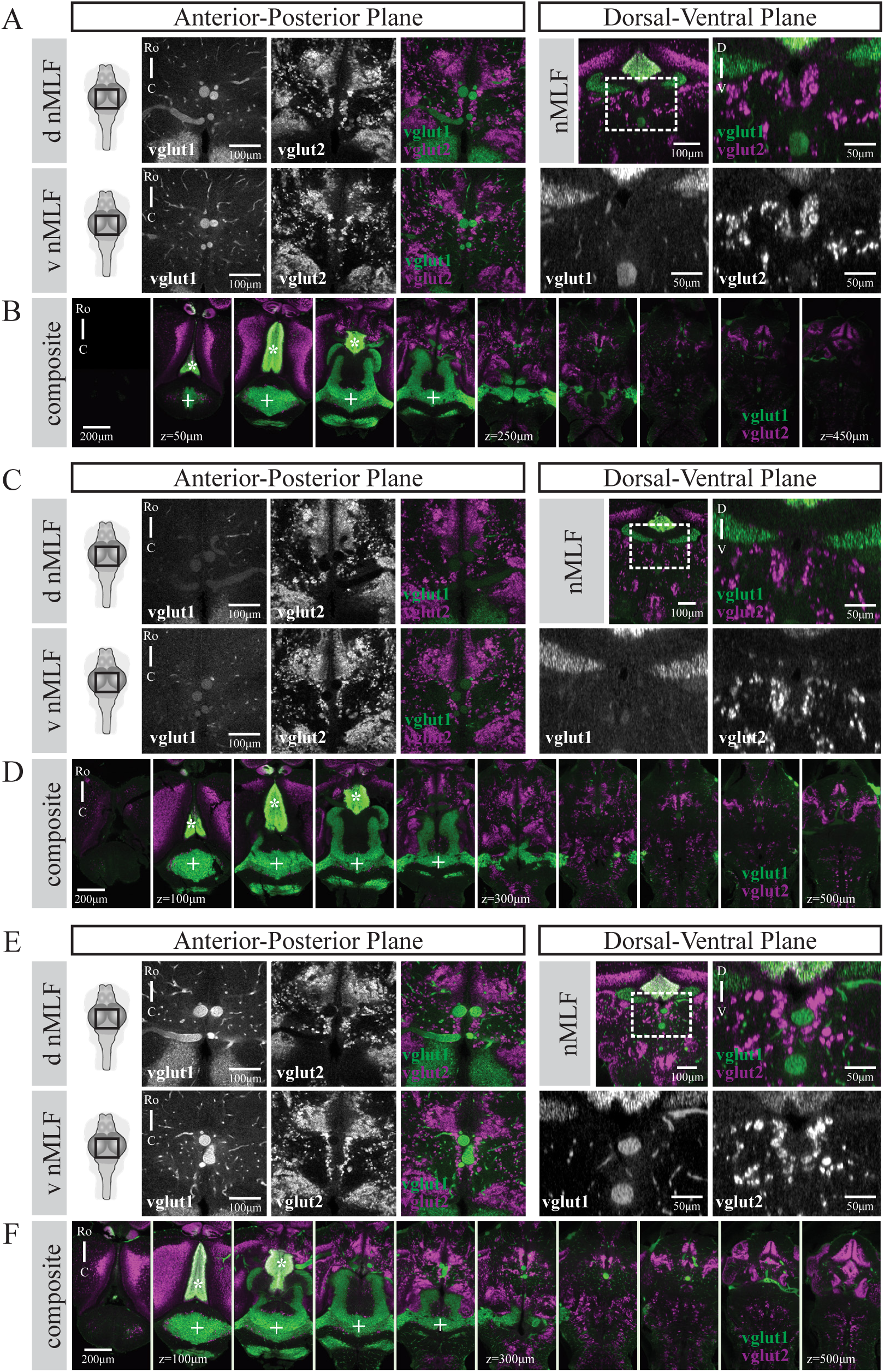
Glutamatergic expression patterns in three 4 week-old juvenile fish (of n=4). A,C,E) Left panels show *vglut1*, *vglut2*, composite in two planes from the dorsal view, scale bar is 100*µ*m. Right panels show *vglut1*, *vglut2*, composite from the transverse view, scale bar is 50*µ*m. B,D,F) Composite images of several planes from dorsal to ventral. Note the presence of torus longitudinalis (*) and cerebellum (+) in *vglut1* (green), with brain wide expression in *vglut2* (magenta). Auto-fluorescence of blood vessels is seen in the *vglut1* channel, as indicated by blue triangles. Scale bar 200*µ*m.

**Figure 6-1:**
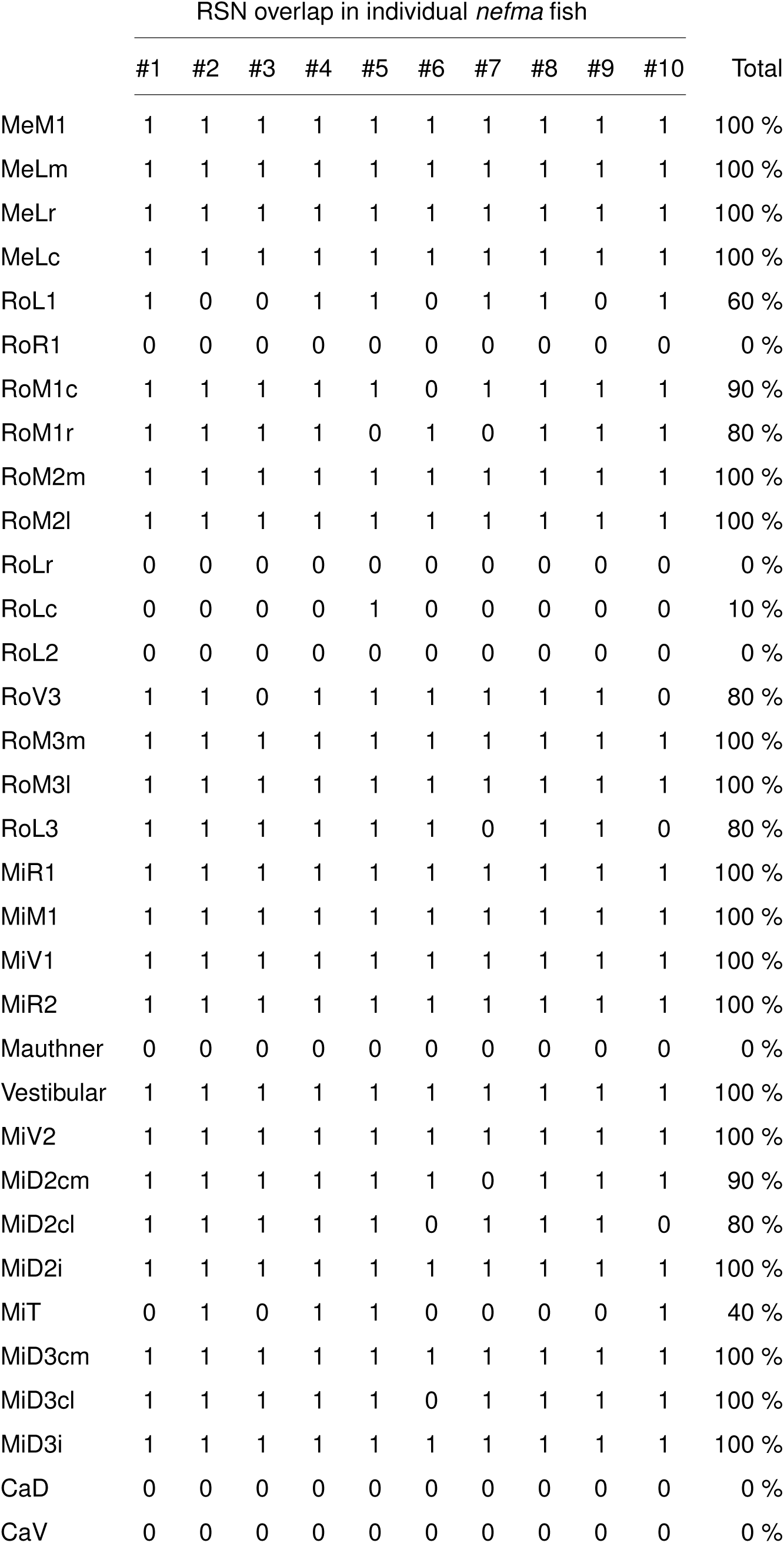
Reticulospinal cells labelled in each *nefma* fish at 6dpf.

**Figure 6-2:**
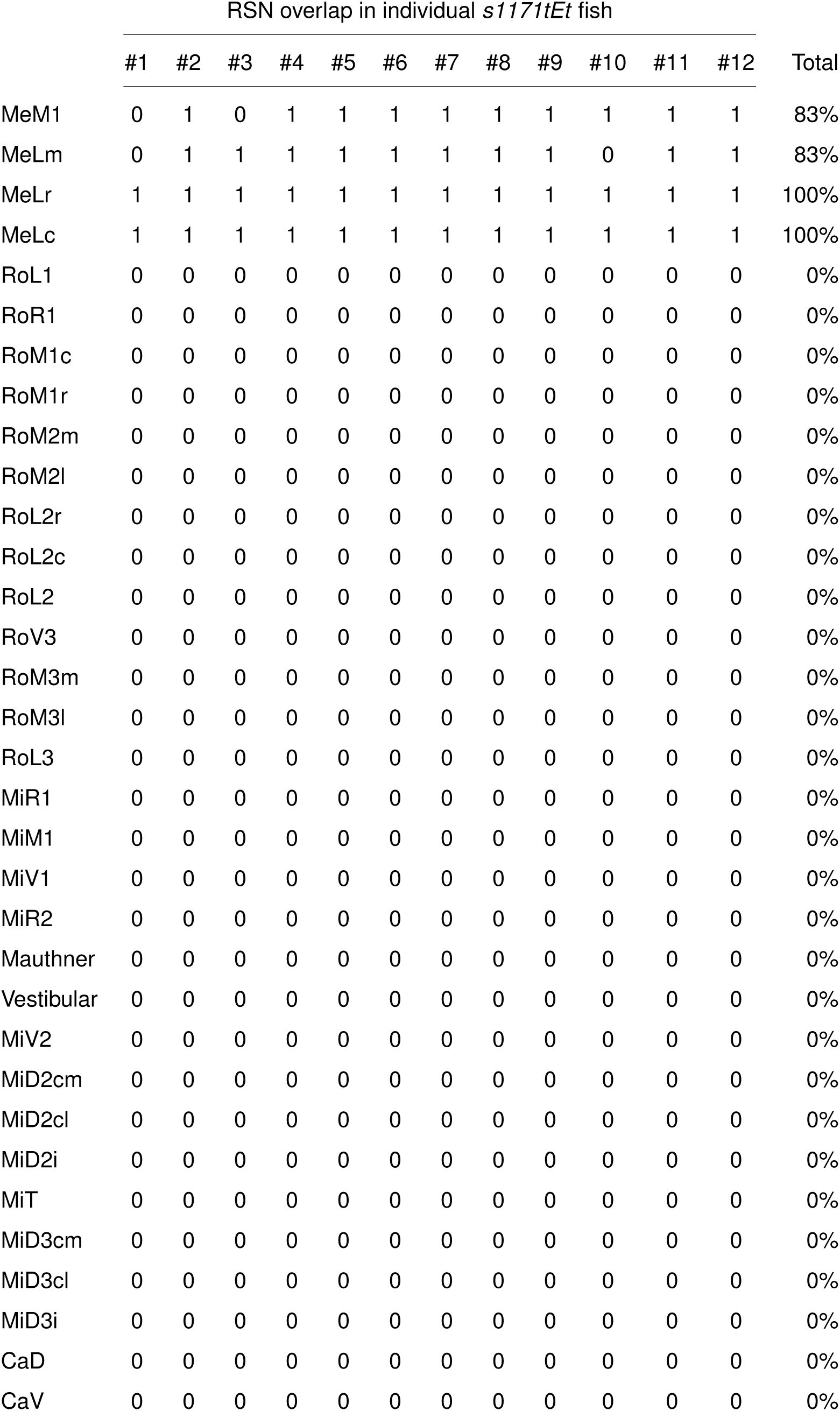
Reticulospinal cells labelled in each *s1171tEt* fish at 6dpf.

**Figure 6-3:**
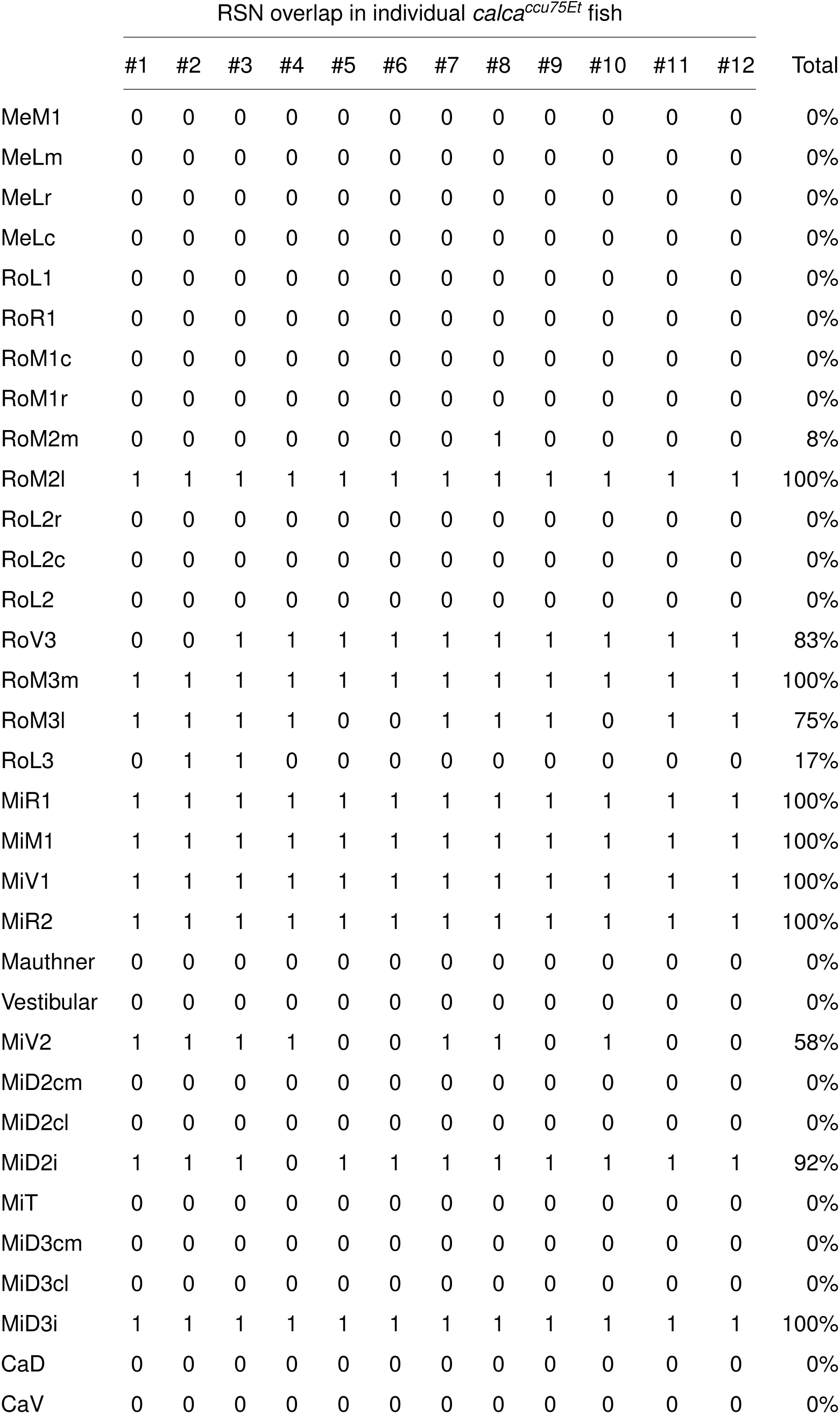
Reticulospinal cells labelled in each *calca^ccu75Et^* fish at 6dpf.

**Figure 6-4:**
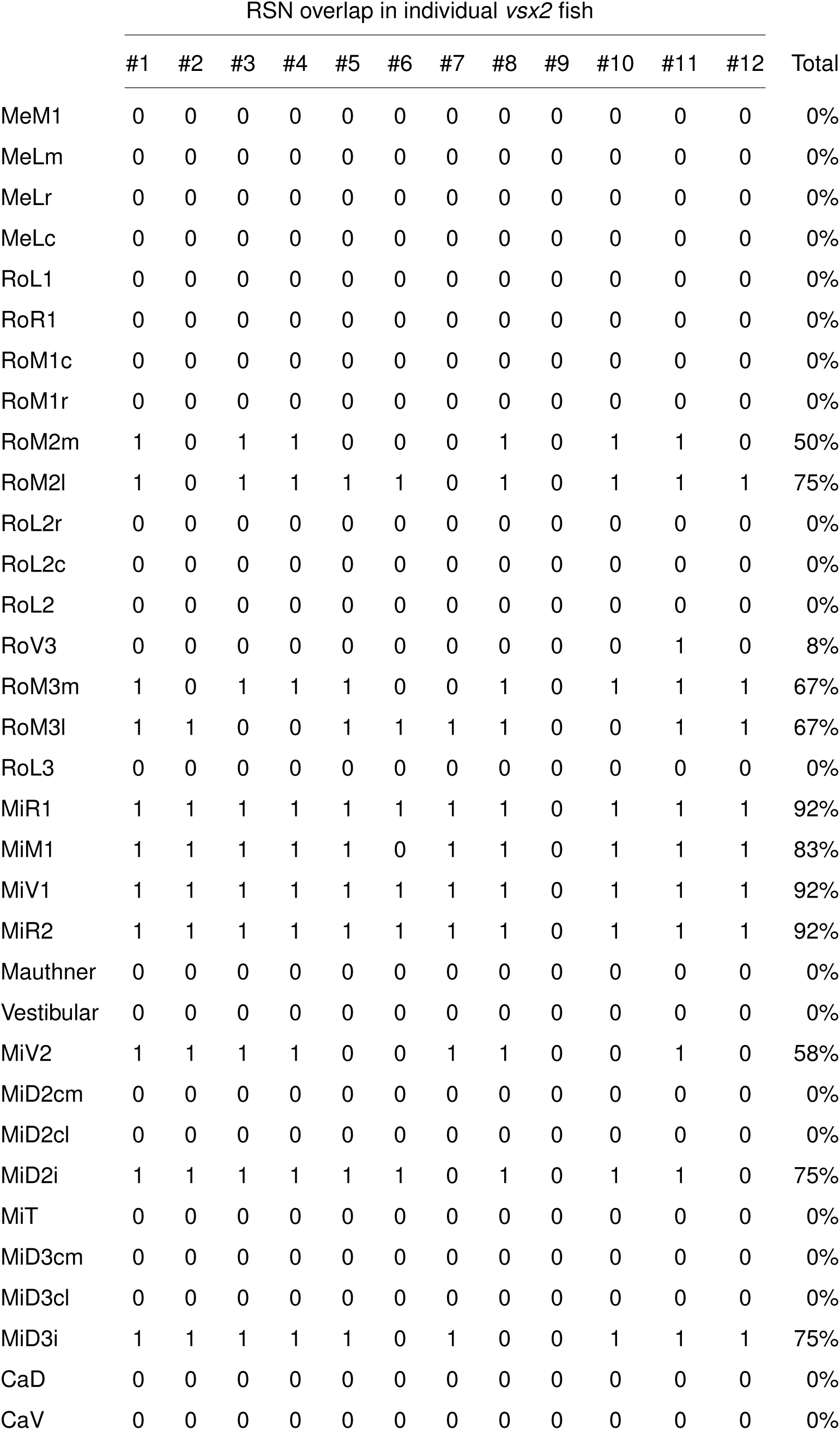
Reticulospinal cells labelled in each *vsx2* fish at 6dpf.

**Figure 6-5:**
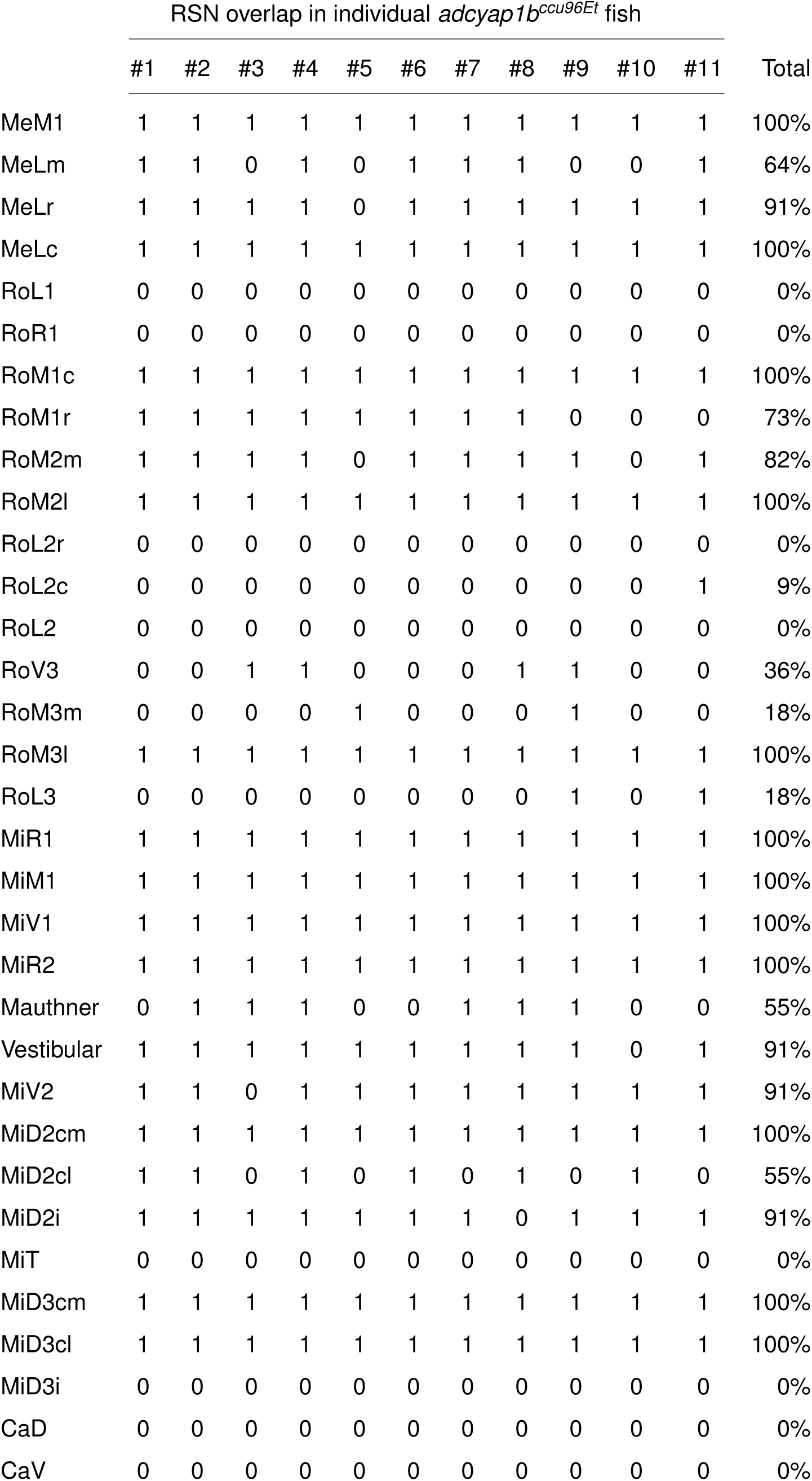
Reticulospinal cells labelled in each *adcyap1b^ccu96Et^* fish at 6dpf.

**Figure 6-6:**
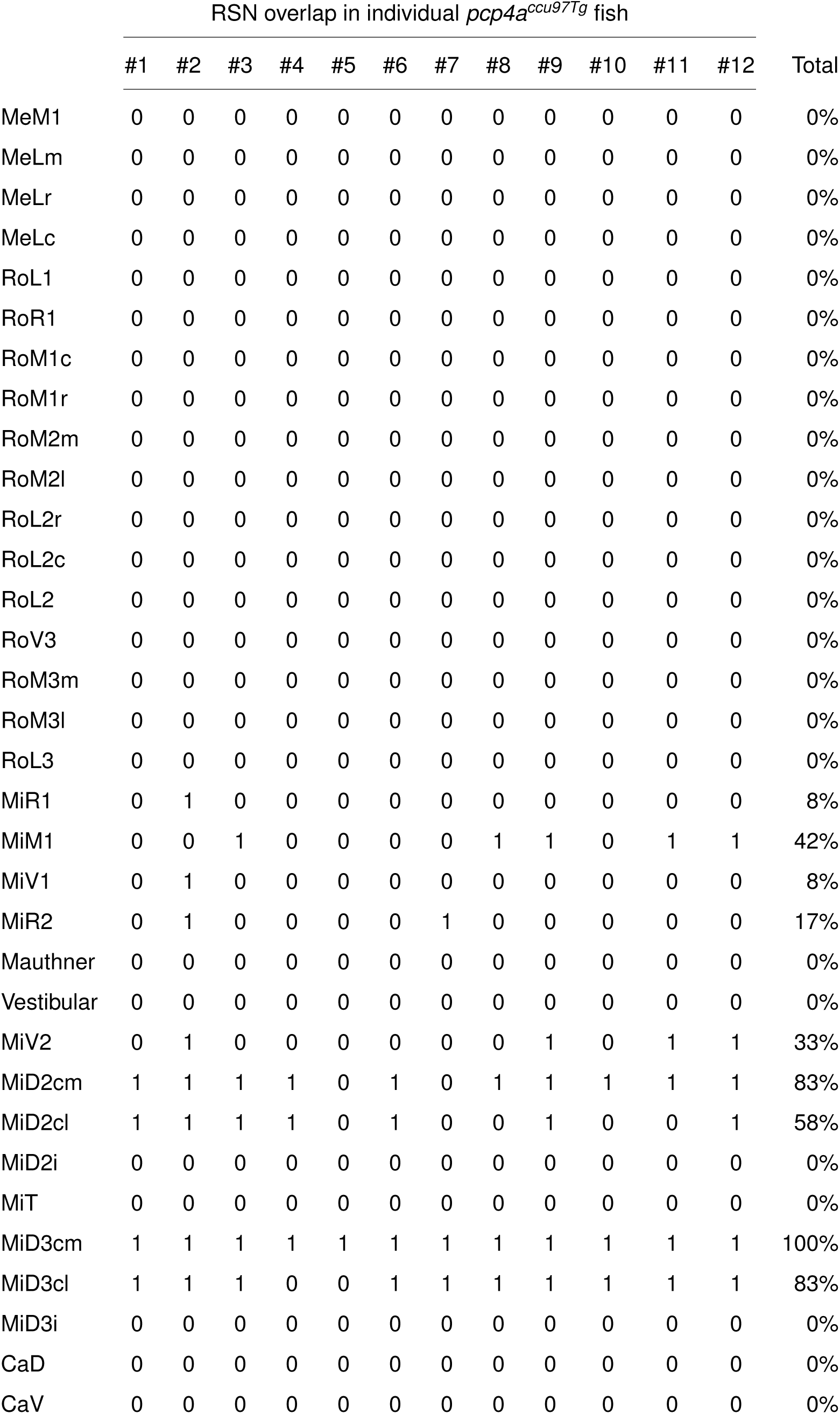
Reticulospinal cells labelled in each *pcp4a^ccu97Tg^* fish at 6dpf.

**Figure 6-7:**
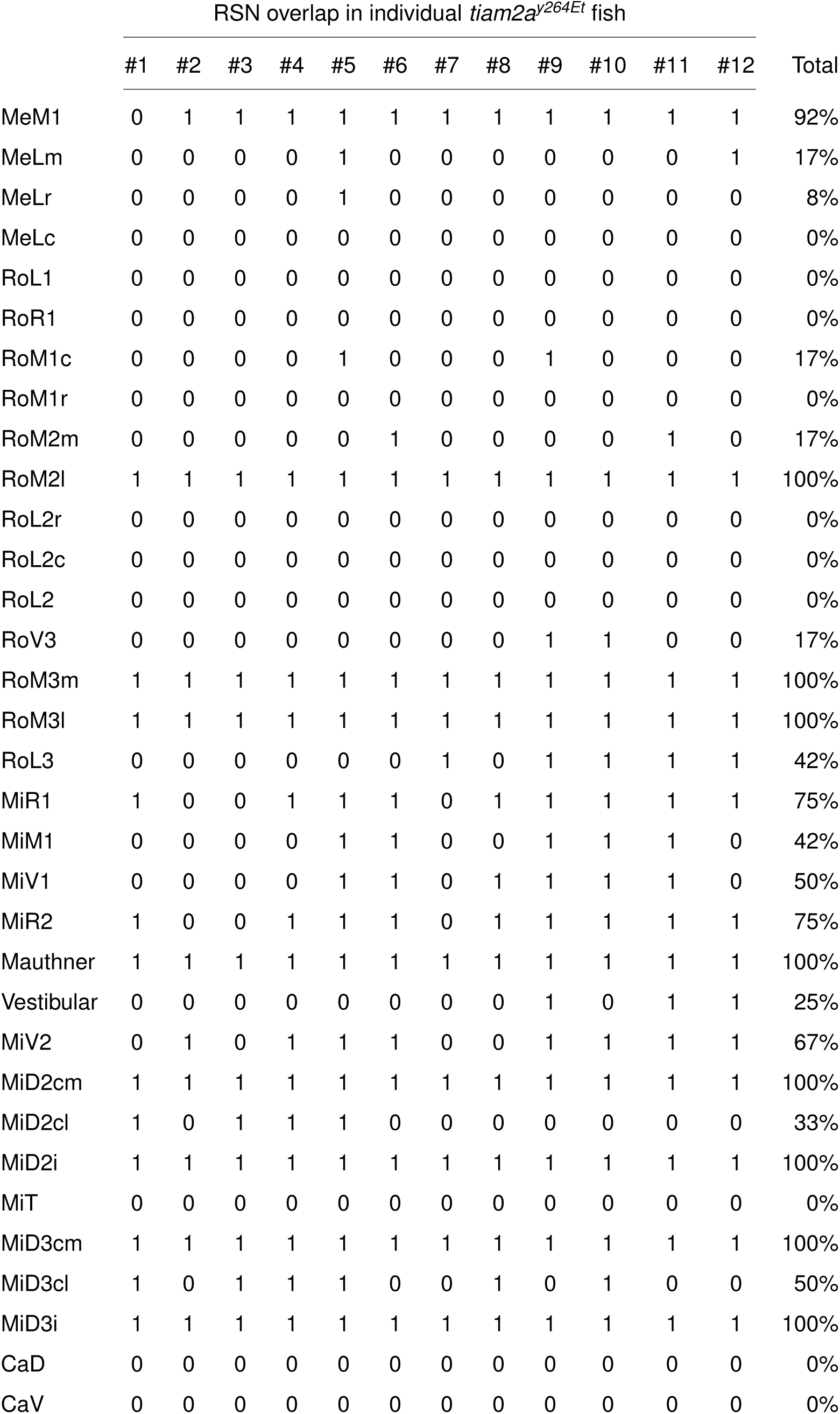
Reticulospinal cells labelled in each *tiam2a^y264Et^* fish at 6dpf.

## References

Akitake CM, Macurak M, Halpern ME, Goll MG (2011) Transgenerational analysis of transcriptional silencing in zebrafish. Developmental Biology 352:191–201.

Alexandre D, Alonzeau J, Bill BR, Ekker SC, Waschek JA (2011) Expression Analysis of PAC1-R and PACAP Genes in Zebrafish Embryos. Journal of Molecular Neuroscience 43:94–100.

Arrenberg AB, Del Bene F, Baier H (2009) Optical control of zebrafish behavior with halorhodopsin. Proceedings of the National Academy of Sciences 106:17968–17973.

Asakawa K, Suster ML, Mizusawa K, Nagayoshi S, Kotani T, Urasaki A, Kishimoto Y, Hibi M, Kawakami K (2008) Genetic dissection of neural circuits by *Tol2* transposon-mediated Gal4 gene and enhancer trapping in zebrafish. Proceedings of the National Academy of Sciences 105:1255–1260.

Avants B, Tustison N, Song G (2009) Advanced normalization tools (ANTS).

Bae YK, Kani S, Shimizu T, Tanabe K, Nojima H, Kimura Y, Higashijima Si, Hibi M (2009) Anatomy of zebrafish cerebellum and screen for mutations affecting its development. Developmental Biology 330:406–426.

Baraban M, Anselme I, Schneider-Maunoury S, Giudicelli F (2013) Zebrafish Embryonic Neurons Transport Messenger RNA to Axons and Growth Cones *In Vivo*. The Journal of Neuroscience 33:15726–15734.

Berg EM, Mrowka L, Bertuzzi M, Madrid D, Picton LD, El Manira A (2023) Brainstem circuits encoding start, speed, and duration of swimming in adult zebrafish. Neuron 111:372–386.e4.

Brownstone RM, Chopek JW (2018) Reticulospinal Systems for Tuning Motor Commands. Frontiers in Neural Circuits 12:30.

Carbo-Tano M, Lapoix M, Jia X, Thouvenin O, Pascucci M, Auclair F, Quan FB, Albadri S, Aguda V, Farouj Y, Hillman EMC, Portugues R, Del Bene F, Thiele TR, Dubuc R, Wyart C (2023) The mesencephalic locomotor region recruits V2a reticulospinal neurons to drive forward locomotion in larval zebrafish. Nature Neuroscience.

Choi HMT, Schwarzkopf M, Fornace ME, Acharya A, Artavanis G, Stegmaier J, Cunha A, Pierce NA (2018) Third-generation *in situ* hybridization chain reaction: multiplexed, quantitative, sensitive, versatile, robust. Development 145:dev165753.

Dal Maschio M, Donovan JC, Helmbrecht TO, Baier H (2017) Linking Neurons to Network Function and Behavior by Two-Photon Holographic Optogenetics and Volumetric Imaging. Neuron 94:774–789.e5.

Datta SR, Patterson GH (2012) Optical highlighter molecules in neurobiology. Current Opinion in Neurobiology 22:111–120.

Distel M, Wullimann MF, Köster RW (2009) Optimized Gal4 genetics for permanent gene expression mapping in zebrafish. Proceedings of the National Academy of Sciences 106:13365–13370.

Eschstruth A, Schneider-Maunoury S, Giudicelli F (2020) Creation of zebrafish knock-in reporter lines in the *nefma* gene by Cas9-mediated homologous recombination. genesis 58.

Fetcho JR, Liu KS (1998) Zebrafish as a Model System for Studying Neuronal Circuits and Behavior. Annals of the New York Academy of Sciences 860:333–345.

Félix R, Markov DA, Renninger SL, Tomás AR, Laborde A, Carey MR, Orger MB, Portugues R (2024) Structural and functional organization of visual responses in the inferior olive of larval zebrafish. The Journal of Neuroscience p. e2352212023.

Gahtan E, O’Malley DM (2001) Rapid lesioning of large numbers of identified vertebrate neurons: applications in zebrafish. Journal of Neuroscience Methods 108:97–110.

Grillner S, El Manira A (2020) Current Principles of Motor Control, with Special Reference to Vertebrate Locomotion. Physiological Reviews 100:271–320.

Hamling KR, Zhu Y, Auer F, Schoppik D (2023) Tilt in Place Microscopy: a Simple, Low-Cost Solution to Image Neural Responses to Body Rotations. The Journal of Neuroscience 43:936–948.

Higashijima Si, Nose A, Eguchi G, Hotta Y, Okamoto H (1997) Mindin/F-spondin Family: Novel ECM Proteins Expressed in the Zebrafish Embryonic Axis. Developmental Biology 192:211–227.

Higashijima S, Mandel G, Fetcho JR (2004) Distribution of prospective glutamatergic, glycinergic, and GABAergic neurons in embryonic and larval zebrafish. Journal of Comparative Neurology 480:1–18.

Josset N, Roussel M, Lemieux M, Lafrance-Zoubga D, Rastqar A, Bretzner F (2018) Distinct Contributions of Mesencephalic Locomotor Region Nuclei to Locomotor Control in the Freely Behaving Mouse. Current Biology 28:884–901.e3.

Kawakami K, Shima A, Kawakami N (2000) Identification of a functional transposase of the *Tol2* element, an *Ac* -like element from the Japanese medaka fish, and its transposition in the zebrafish germ lineage. Proceedings of the National Academy of Sciences 97:11403–11408.

Kenney JW, Steadman PE, Young O, Shi MT, Polanco M, Dubaishi S, Covert K, Mueller T, Frankland PW (2021) A 3D adult zebrafish brain atlas (AZBA) for the digital age. eLife 10:e69988.

Kimmel CB, Hatta K, Metcalfe WK (1990) Early axonal contacts during development of an identified dendrite in the brain of the zebrafish. Neuron 4:535–545.

Kimmel CB, Powell SL, Metcalfe WK (1982) Brain neurons which project to the spinal cord in young larvae of the zebrafish. Journal of Comparative Neurology 205:112–127.

Kimura Y, Okamura Y, Higashijima Si (2006) *alx*, a Zebrafish Homolog of *Chx10*, Marks Ipsilateral Descending Excitatory Interneurons That Participate in the Regulation of Spinal Locomotor Circuits. The Journal of Neuroscience 26:5684–5697.

Kimura Y, Satou C, Fujioka S, Shoji W, Umeda K, Ishizuka T, Yawo H, Higashijima Si (2013) Hindbrain V2a Neurons in the Excitation of Spinal Locomotor Circuits during Zebrafish Swimming. Current Biology 23:843–849.

Kinkhabwala A, Riley M, Koyama M, Monen J, Satou C, Kimura Y, Higashijima Si, Fetcho J (2011) A structural and functional ground plan for neurons in the hindbrain of zebrafish. Proceedings of the National Academy of Sciences 108:1164–1169.

Kunst M, Laurell E, Mokayes N, Kramer A, Kubo F, Fernandes AM, Förster D, Dal Maschio M, Baier H (2019) A Cellular-Resolution Atlas of the Larval Zebrafish Brain. Neuron 103:21–38.e5.

Kwan KM, Fujimoto E, Grabher C, Mangum BD, Hardy ME, Campbell DS, Parant JM, Yost HJ, Kanki JP, Chien C (2007) The Tol2kit: A multisite gateway-based construction kit for *Tol2* transposon transgenesis constructs. Developmental Dynamics 236:3088–3099.

Le Ray D, Bertrand SS, Dubuc R (2022) Cholinergic Modulation of Locomotor Circuits in Vertebrates. International Journal of Molecular Sciences 23:10738.

Liu Z, Bagnall MW (2023) Organization of vestibular circuits for postural control in zebrafish. Current Opinion in Neurobiology 82:102776.

Liu Z, Kimura Y, Higashijima Si, Hildebrand DG, Morgan JL, Bagnall MW (2020) Central Vestibular Tuning Arises from Patterned Convergence of Otolith Afferents. Neuron 108:748–762.e4.

Marquart GD, Tabor KM, Brown M, Strykowski JL, Varshney GK, LaFave MC, Mueller T, Burgess SM, Higashijima Si, Burgess HA (2015) A 3D Searchable Database of Transgenic Zebrafish Gal4 and Cre Lines for Functional Neuroanatomy Studies. Frontiers in Neural Circuits 9.

Martins S, Monteiro JF, Vito M, Weintraub D, Almeida J, Certal AC (2016) Toward an Integrated Zebrafish Health Management Program Supporting Cancer and Neuroscience Research. Zebrafish 13:S–47–S–55.

McLean DL, Fetcho JR (2004) Relationship of tyrosine hydroxylase and serotonin immunoreactivity to sensorimotor circuitry in larval zebrafish. Journal of Comparative Neurology 480:57–71.

Metcalfe WK, Mendelson B, Kimmel CB (1986) Segmental homologies among reticulospinal neurons in the hindbrain of the zebrafish larva. Journal of Comparative Neurology 251:147–159.

Mione M, Lele Z, Kwong CT, Concha ML, Clarke JD (2006) Expression of *pcp4a* in subpopulations of CNS neurons in zebrafish. Journal of Comparative Neurology 495:769–787.

O’Malley DM, Kao YH, Fetcho JR (1996) Imaging the Functional Organization of Zebrafish Hindbrain Segments during Escape Behaviors. Neuron 17:1145–1155.

Orger MB, Kampff AR, Severi KE, Bollmann JH, Engert F (2008) Control of visually guided behavior by distinct populations of spinal projection neurons. Nature Neuroscience 11:327–333.

Perreault MC, Giorgi A (2019) Diversity of reticulospinal systems in mammals. Current Opinion in Physiology 8:161–169.

Randlett O, Wee CL, Naumann EA, Nnaemeka O, Schoppik D, Fitzgerald JE, Portugues R, Lacoste AMB, Riegler C, Engert F, Schier AF (2015) Whole-brain activity mapping onto a zebrafish brain atlas. Nature Methods 12:1039–1046.

Scott EK, Baier H (2009) The cellular architecture of the larval zebrafish tectum, as revealed by gal4 enhancer trap lines. Frontiers in Neural Circuits 3:13.

Semmelhack JL, Donovan JC, Thiele TR, Kuehn E, Laurell E, Baier H (2014) A dedicated visual pathway for prey detection in larval zebrafish. eLife 3:e04878.

Severi K, Portugues R, Marques J, O’Malley D, Orger M, Engert F (2014) Neural Control and Modulation of Swimming Speed in the Larval Zebrafish. Neuron 83:692–707.

Siegel JM (1979) Behavioral functions of the reticular formation. Brain Research Reviews 1:69–105.

Sugioka T, Tanimoto M, Higashijima Si (2023) Biomechanics and neural circuits for vestibular-induced fine postural control in larval zebrafish. Nature Communications 14:1217.

Suster ML, Abe G, Schouw A, Kawakami K (2011) Transposon-mediated BAC transgenesis in zebrafish. Nature Protocols 6:1998–2021.

Tabor KM, Bergeron SA, Horstick EJ, Jordan DC, Aho V, Porkka-Heiskanen T, Haspel G, Burgess HA (2014) Direct activation of the Mauthner cell by electric field pulses drives ultrarapid escape responses. Journal of Neurophysiology 112:834–844.

Tabor KM, Smith TS, Brown M, Bergeron SA, Briggman KL, Burgess HA (2018) Presynaptic Inhibition Selectively Gates Auditory Transmission to the Brainstem Startle Circuit. Current Biology 28:2527–2535.e8.

Tada M, Takeuchi A, Hashizume M, Kitamura K, Kano M (2014) A highly sensitive fluorescent indicator dye for calcium imaging of neural activity in vitro and in vivo. The European Journal of Neuroscience 39:1720–1728.

Teh C, Chudakov DM, Poon KL, Mamedov IZ, Sek JY, Shidlovsky K, Lukyanov S, Korzh V (2010) Optogenetic in vivocell manipulation in KillerRed-expressing zebrafish transgenics. BMC Developmental Biology 10:110.

Thiele T, Donovan J, Baier H (2014) Descending Control of Swim Posture by a Midbrain Nucleus in Zebrafish. Neuron 83:679–691.

Thisse B, Heyer V, Lux A, Alunni V, Degrave A, Seiliez I, Kirchner J, Parkhill JP, Thisse C (2004) Spatial and Temporal Expression of the Zebrafish Genome by Large-Scale In Situ Hybridization Screening In Methods in Cell Biology, Vol. 77, pp. 505–519. Elsevier.

Van Opbergen CJ, Koopman CD, Kok BJ, Knöpfel T, Renninger SL, Orger MB, Vos MA, Van Veen TA, Bakkers J, De Boer TP (2018) Optogenetic sensors in the zebrafish heart: a novel in vivo electrophysiological tool to study cardiac arrhythmogenesis. Theranostics 8:4750–4764.

Woods IG, Schoppik D, Shi VJ, Zimmerman S, Coleman HA, Greenwood J, Soucy ER, Schier AF (2014) Neuropeptidergic Signaling Partitions Arousal Behaviors in Zebrafish. The Journal of Neuroscience 34:3142–3160.

Wu S, Adams BA, Fradinger EA, Sherwood NM (2006) Role of Two Genes Encoding PACAP in Early Brain Development in Zebrafish. Annals of the New York Academy of Sciences 1070:602–621.

Wullimann MF, Rupp B, Reichert H (1996) Neuroanatomy of the Zebrafish Brain Birkhäuser Basel, Basel.

Wyart C, Bene FD, Warp E, Scott EK, Trauner D, Baier H, Isacoff EY (2009) Optogenetic dissection of a behavioural module in the vertebrate spinal cord. Nature 461:407–410.

Yamamoto N, Nakayama T, Hagio H (2017) Descending pathways to the spinal cord in teleosts in comparison with mammals, with special attention to rubrospinal pathways. *Development*, Growth & Differentiation 59:188–193.

Zhang Y, Ouyang J, Qie J, Zhang G, Liu L, Yang P (2019) Optimization of the Gal4/UAS transgenic tools in zebrafish. Applied Microbiology and Biotechnology 103:1789–1799.

Zhu P, Fajardo O, Shum J, Zhang Schärer YP, Friedrich RW (2012) High-resolution optical control of spatiotemporal neuronal activity patterns in zebrafish using a digital micromirror device. Nature Protocols 7:1410–1425.

